# Systemic and intrinsic functions of ATRX in glial cell fate and CNS myelination

**DOI:** 10.1101/2022.09.15.508143

**Authors:** Megan E. Rowland, Yan Jiang, Sarfraz Shafiq, Alireza Ghahramani, Miguel A. Pena-Ortiz, Vanessa Dumeaux, Nathalie G. Bérubé

## Abstract

Neurodevelopmental disorders are often characterized by abnormal production of myelin, an extension of the oligodendrocyte plasma membrane wrapped around axons to facilitate nerve conduction. However, the molecular mechanisms that control myelination during brain development are incompletely resolved. Here, we provide evidence that loss of ATRX, encoded by the gene mutated in the ATR-X intellectual disability syndrome, leads to myelin deficits in the mouse CNS. While postnatal systemic thyroxine administration can improve myelination, the rescue is incomplete, pointing to additional roles of ATRX in this process. We show that targeted inactivation of ATRX in postnatal oligodendrocyte progenitor cells (OPCs), but not in neurons, also leads to myelination deficits, demonstrating cell-intrinsic effects of ATRX deficiency. A subset of ATRX-null OPCs express lower levels of oligodendrocyte specification and differentiation markers, including the basic helix-loop-helix Olig2 transcription factor. Mechanistically, we provide evidence that ATRX occupies genomic sites in OPCs marked by H3K27Ac, CHD7 and CHD8 and demonstrate that reduced *Olig2* expression is associated with decreased H3K27Ac. Finally, our data suggest that ATRX-null OPCs acquire a more plastic state and can exhibit astrocyte-like features *in vitro* and *in vivo*, supporting a model in which ATRX regulates the onset of myelination by promoting OPC identity and suppressing astrogliogenesis. These previously unrecognized functions of ATRX might explain white matter pathogenesis in ATR-X syndrome patients.

## Introduction

Myelin is an extension of the oligodendrocyte (OL) plasma membrane that wraps itself around axons to increase conduction velocity, enabling efficient propagation of action potentials. A series of events are necessary to produce myelin, including proliferation and migration of oligodendrocyte precursor cells (OPCs), recognition of recipient axons, membrane expansion, wrapping of axons and the appropriate production of membrane components and myelin proteins for compaction (*1*).

In the developing neocortex, the generation of neurons from neural progenitor cells eventually shifts towards production of astrocytes and then OLs (*2*). The majority of primitive OPCs are generated postnatally from neural stem cells (NSCs) and intermediate glial progenitor cells (iGCs) through the expression of the oligodendrocyte basic-helix-loop-helix transcription factors Olig1 and Olig2 (*3*). Olig2 is required for OPC generation and suppresses the astrocytic lineage (*4–7*). Downstream of Olig2, the SRY-box transcription factor 10 (Sox10) participates in a positive feedback loop to establish the OPC lineage (*8–10*). Notably, Sox10 activates several OPC-critical genes, including *Chondroitin sulfate proteoglycan* (*Cspg4*; protein: neural glial antigen 2 (NG2)) and *Platelet-derived growth factor receptor α* (*Pdgfrα*) (*11–13*). NG2 is a surface type I transmembrane core protein involved in cell migration and proliferation (*14*), whereas PDGFRα promotes OPC proliferation, migration and survival (*15, 16*). Further commitment to the OL lineage starts with the generation of immature oligodendrocytes (iOLs) which, after cell cycle exit, express myelin regulatory factor (MRF) (*17*). As iOLs mature and contact axons, they become highly branched and upregulate several myelin sheath proteins in order to encompass the axon (*1, 18*). The expression of myelin proteins is complex and regulated by several different transcription factors including the aforementioned Olig1 (*19*), Sox10 (*19, 20*) and MRF (*17*) but also Sp1 (*21*), Nkx2.2 (*22*) and the ligand-bound thyroid hormone receptor (*23–25*).

Myelination has been studied extensively in the context of demyelinating diseases like multiple sclerosis and leukodystrophies (*26*). However, myelin deficits are also frequently observed in neurodevelopmental cognitive disorders like autism and schizophrenia and can contribute to intellectual disability (*27–29*). This is exemplified by the ATR-X syndrome, an intellectual disability disorder caused by abnormalities in the ATRX chromatin remodeling protein (*30*). Affected males often exhibit cognitive dysfunction, seizures and autistic features, among other developmental defects (*31–35*). ATR-X syndrome patients also display abnormal myelination with a prevalence of 44% (*33, 34*) making white matter abnormalities a supporting feature of the diagnosis.

ATRX belongs to the Switch/Sucrose non-fermenting (Swi/Snf) family of chromatin remodelers (*36*), and can regulate gene expression through several mechanisms. In the newborn cortex, ATRX represses expression of imprinted genes by fostering long-range chromatin interactions mediated by CCCTC-binding factor (CTCF) and cohesin (*37, 38*). In embryonic stem cells, it promotes maintenance of the repressive H3K27me3 histone mark at polycomb target genes (*39*). On the other hand, ATRX can promote gene transcription through the incorporation of histone H3.3 and facilitating RNA PolII elongation through G-rich sequences (*40, 41*). In testes, ATRX regulates imprinted genes (*42*) associates with the androgen receptor and facilitates transcription of its target genes (*43*). ATRX also promotes the expression of α-globin by negatively regulating the histone variant macroH2A (*44*).

The molecular and cellular basis of white matter abnormalities associated with ATRX dysfunction has not yet been investigated. Here, we report that deletion of *Atrx* in the central nervous system causes hypomyelination. While myelin levels and OL numbers were partially recovered upon administration of the thyroid hormone thyroxine (T4), this treatment failed to replenish OPCs, indicating that ATRX is required to maintain these progenitors independently of circulating T4 levels. Indeed, targeted inactivation of *Atrx* in Sox10-expressing OPCs resulted in hypomyelination, revealing cell autonomous functions of ATRX in OPCs. We show that ATRX occupies and maintains *Olig2* gene expression. In the absence of ATRX, OPCs tend to revert to a more malleable state, and gain the ability to differentiate into astrocyte-like cells. We conclude that ATRX deficiency affects the developmental trajectory of glial precursors, which could be the underlying cause of defective myelination in the associated ATR-X syndrome.

## Materials and Methods

### Animal husbandry and genotyping

Mice were exposed to 12-hour light/12-hour dark cycles and fed water and regular chow ad libitum. The *Atrx*^loxP^ mice have been described previously (*45*) (MGI:3528480). Briefly, *loxP* sites flanking exon 18 of the *Atrx* gene allow for deletion of this exon, destabilization of the mRNA and absence of the full-length ATRX protein (*45*). Mating *Atrx*^loxP^ female mice to males expressing Cre recombinase under the control of the *Foxg1* promoter (129(Cg)-*Foxg1^tm1(cre)Skm^*/J, RRID:IMSR_JAX:004337) (*46*) yielded male progeny with ATRX deficiency in the forebrain and anterior pituitary (*Atrx*^loxP^;FoxG1Cre RRID:MGI:3530074) called *Atrx*^FoxG1Cre^ for simplicity. Mating *Atrx*^loxP^ female mice to males expressing Cre recombinase under the control of the *Nex* gene promoter (*47, 48*) (Neurod6^tm1(cre)Kan^, MGI:2668659) produced male progeny with ATRX deficiency in forebrain glutamatergic neurons (*Atrx*^loxP^;NexCre or *Atrx*^NexCre^ for simplicity). Mating *Atrx*^loxP^ female mice to males expressing Cre recombinase under the control of the inducible Sox10 promoter (*49*) (CBA;B6-Tg(Sox10-icre/ERT2)388Wdr, MGI:5634390, RRID:IMSR JAX:027651), produced male progeny with *Atrx* deficiency in OPCs (*Atrx*^loxP^;Sox10Cre or *Atrx*^Sox10Cre^) upon tamoxifen treatment. Two Cre-sensitive reporter lines were bred into the *Atrx*^loxP^ mice for nuclei sorting and lineage tracing. The Sun1GFP allele (B6;*129-Gt(ROSA)26Sor^tm5(CAG-Sun1/sfGFP)Nat^*/J, MGI:5614796, RRID: IMSR_JAX:021039) encodes the Sun1GFP fusion protein (located in the nuclear membrane) upon Cre-mediated recombination (*50*) and the Tomato-Ai14 (Ai14) allele (B6.Cg-*Gt(ROSA)26Sor^tm14(CAG-tdTomato)Hze^*/J, MGI:3809524, RRID:IMSR_JAX:007914) (*51*) expresses the tdTomato protein with red fluorescence in the cytoplasm and nucleus upon Cre recombination. Ear punch genomic DNA was used for PCR genotyping of the *Atrx* floxed or wildtype alleles using the primers 17F, 18R and neoR as described previously (*45*). The *FoxG1Cre, NexCre, Sox10Cre, Tomato-Ai14* and *Sun1GFP* genotyping primers are listed in **Supplementary Table 1**.

### Microarray analysis

Total forebrain RNA (10 μg) was isolated from three pairs of littermate matched *Atrx*^FoxG1Cre^ and control P17 mice using the RNeasy Mini kit (Qiagen Cat# 74104). cRNA was generated and hybridized to an Affymetrix Mouse Genome 430 2.0 Array at the London Regional Genomics Center (London, Canada). Probe signal intensities were generated using GCOS1.4 (Affymetrix Inc., Santa Clara, CA) using default values for the Statistical Expression algorithm parameters and a Target Signal of 150 for all probe sets and a Normalization Value of 1. Gene level data was generated using the RNA preprocessor in GeneSpring GX 7.3.1 (Agilent Technologies Inc., Palo Alto, CA). Data were then transformed (measurements less than 0.01 set to 0.01), normalized per chip to the 50th percentile, and per gene to control samples. Probe sets representing *Atrx* transcripts were removed (10 sets). The remaining probe sets were filtered by fold change ≥1.5 between control and *Atrx*^FoxG1Cre^ samples, and by confidence level of p<0.05. Significantly overrepresented GO categories were determined using the Enrichr web platform (*52–54*).

### Thyroxine and tamoxifen injections

Subcutaneous injection of L-thyroxine (0.1 mg/kg; Sigma Cat# T2376-100MG) was performed daily on control and *Atrx*^Foxg1Cre^ mice from birth (P0) until P14 as done previously (*55*). Tamoxifen (10mg; Cat# T5648, Sigma) was dissolved in 100 μL 95% ethanol at 65°C for 10 mins, followed by dilution in 900 μL corn oil (Cat# C8267, Sigma). Lactating mothers were injected intraperitoneally daily with 2 mg tamoxifen for two (glial culture experiments) or three consecutive days (*56*).

### Western blot analysis

Protein was extracted from P14 or P20 mouse forebrain and homogenized in RIPA buffer (150 mM NaCl, 1% NP-40, 50 mM Tris pH 8.0, 0.5% deoxycholic acid, 0.1% SDS, 0.2 mM PMSF, 0.5 mM NaF, 0.1 mM Na_3_VO_4_, 1x protease inhibitor cocktail (Millipore Sigma Cat# 11873580001). Homogenized tissue was incubated on ice for 20 minutes and centrifuged for 20 minutes at 1×10^4^ RPM. Supernatant protein was quantified using the Bradford assay (BioRad Cat# 500-0006). Protein (30 μg) was resolved on a 12% SDS-PAGE and transferred to a 0.45 μm nitrocellulose membrane (BioRad Cat# 1620115). The membranes were probed with anti-MAG, mouse monoclonal (1:3000, Abcam Cat# ab89780, RRID:AB_2042411), anti-MOG, rabbit polyclonal (1:3000, Abcam Cat# ab32760, RRID:AB_2145529), or anti-MBP, rat monoclonal (1:3000, Abcam Cat# ab7349, RRID:AB_305869) antibodies at 4°C overnight. This was followed by incubation with the appropriate horseradish peroxidase–conjugated secondary antibody for 1 hour at room temperature: mouse anti-HRP (1:3,000, Santa Cruz Cat# sc-516102, RRID:AB_2687626), rabbit anti-HRP (1:5000, Jackson ImmunoResearch Cat# 111-036-003, RRID:AB_2337942) or rat anti-HRP (1:3,000, Santa Cruz Cat# sc-2006, RRID:AB_1125219). The membrane was incubated in enhanced chemiluminescent solution (Thermo Fisher Cat# 34095) before exposure using film (Progene Cat# 39-20810) or Universal Hood III (BioRad Cat# 731BR00882) and analyzed with Image Lab (BioRad, Version 4.1, 2012).

### Mixed glial primary culture

Cre recombination in control and *Atrx*^Sox10Cre^ pups was induced by IP injection of 2 mg tamoxifen to lactating mothers for two consecutive days (P0-P1) (*56*). At P3, mouse cortices were dissected into MEM and tissue dissociated by pipetting with a P1000. Samples were then incubated at 37°C with OPC papain solution (Worthington Cat#LS003124) for 20 minutes with constant inversion to prevent tissue aggregation. Papain was deactivated with serum-supplemented media for 10 min at room temperature followed by trituration with a flame polished Pasteur pipette. Samples were centrifuged at 300 g for 5 minutes and pellets resuspended in 1 mL DMEM supplemented with 10% FBS (Gibco Cat # 12483020), 1% Antibiotic-Antimycotic (Gibco Cat # 15240062) and 1% GlutaMax (Gibco Cat # 35050061). The cell suspension from each tube was added to a pre-equilibrated poly-L-lysine (1 mg/mL) coated 6 well plate in 1 mL of media or flasks with 5 mL of media. Cultures were incubated in the presence of 8.5% CO_2_ in a tissue culture incubator for 3 hours to allow cells to attach to the poly-L-lysine substrate followed by a full media change. A 2/3 volume media change was performed every three days (*57, 58*).

### Immunofluorescence

P14 or P20 mice were trans-cardially perfused with PBS followed by 4% paraformaldehyde. Brains were fixed in 4% paraformaldehyde overnight. The next day, brains were washed 3 times for 5 min in PBS, sunk in 30% sucrose, flash-frozen on dry ice in Cryomatrix cryoprotectant (ThermoFisher Cat# 6769006), sectioned coronally at 8 μm thickness (Leica CM 3050S) on Superfrost slides (Thermo Fisher Cat# 22-037-246) and stored at −80°C with a desiccant (VWR, 61161-319). For immunofluorescence staining, slides were rehydrated in 1x PBS for 5 minutes, washed with PBS + 0.1% TritonX-100 (Millipore Sigma Cat# T8787), blocked with 10% normal goat serum diluted in washing solution (Millipore Sigma Cat# G9023) for 1 hour and incubated with primary antibody overnight at 4°C. Slides were washed 3 times for 5 min with PBS and incubated with secondary antibody for 1 hour in the dark, and then washed twice for 5 min in PBS. Sections were counterstained with 1 μg/mL DAPI (Millipore Sigma Cat# D9542) for 5 min, washed for 5 min with PBS and mounted with Permafluor (Thermo Fisher Cat# TA-006-FM). Primary mixed glial cultures were fixed for immunofluorescence staining at 5, 6 and 9DIV (days *in vitro*) with 3% PFA at room temperature for 15 minutes. Cells were permeabilized with 0.1% TritonX-100/PBS (Millipore Sigma Cat# T8787), blocked with 10% normal goat serum in PBS (Millipore Sigma Cat# G9023) for 1 hour and incubated with primary antibody overnight at 4°C. The next day, flasks were washed 3 times for 5 minutes with PBS and incubated with the secondary antibody diluted in blocking solution for 1 hour at room temperature. Cells were then counter-stained with 1 μg/mL DAPI (Millipore Sigma Cat# D9542) for 5 min, washed for 5 min with PBS and mounted with Permafluor (Thermo Fisher Cat# TA-006-FM). The following primary antibodies were used: anti-MOG, rabbit polyclonal (1:200, Abcam Cat# ab32760, RRID:AB_2145529), anti-MBP, rat monoclonal (1:50, Abcam Cat# ab7349, RRID:AB_305869), anti-ATRX, rabbit polyclonal (1:75, Santa Cruz Biotechnology Cat# sc-15408, RRID:AB_2061023), anti-Olig2, rabbit polyclonal (1:200, Millipore Cat# AB9610, RRID:AB_570666), anti-S100β, rabbit polyclonal (1:200, Agilent Cat# Z0311, RRID:AB_10013383), anti-GFAP, rabbit polyclonal (1:200, Agilent Cat# Z0334, RRID:AB_10013382), anti-NG2, rabbit polyclonal (1:200, Millipore Cat# AB5320, RRID:AB_11213678), anti-PDGFRα, rabbit polyclonal (1:200 Abcam Cat# ab65258, RRID:AB_1141669), anti-APC, mouse monoclonal (Abcam Cat# ab16794, RRID:AB_443473, anti-Ki67, rabbit polyclonal (1:150 Abcam Cat# ab15580, RRID:AB_443209) and anti-NFIA (1:100 Sigma-Aldrich Cat# HPA006111, RRID:AB_1854422). The secondary antibodies used were goat anti-rabbit-Alexa Fluor 594 (1:800, Thermo Fisher Scientific, A-11012, RRID:AB_2534079), goat anti-rabbit-Alexa Fluor 488 (1:800, Thermo Fisher Scientific Cat# A-11008, RRID:AB_143165), goat anti-mouse-Alexa Fluor 594 (1:800 Thermo Fisher Scientific Cat# A-21125, RRID:AB_2535767), goat anti-mouse-Alexa Fluor 488 (1:800, Thermo Fisher Scientific Cat# A-11001, RRID:AB_2534069) and goat anti-rat-Alexa Fluor 488 (1:800, Thermo Fisher Scientific, A-11006, RRID:AB_2534074). In Situ Cell Death Detection kit (Roche Cat# 11684795910) was used to assess cell death.

### Fluorescence microscopy, imaging and cell counts

Immunofluorescence images were captured using an inverted microscope (DMI 6000b, Leica) outfitted with a digital camera (ORCA-ER, Hamamatsu). Openlab software (PerkinElmer Version 5.0, RRID:SCR_012158) was used for image capture. Image processing was performed using Volocity (PerkinElmer Demo Version 6.0.1, RRID:SCR_002668) and Adobe Photoshop. Cell counts of brain sections and cultured cells were performed in a blinded and randomized manner in control or experimental samples using Volocity (PerkinElmer Demo Version 6.0.1, RRID:SCR_002668). For brain sections, counts were performed in 2-10 sections per biological replicate and 3-4 biological replicates were counted for each genotype.

### Fluorescence activated sorting of OPC nuclei

Fluorescence-activated nuclei sorting (FANS) was used to purify Sun1-GFP^+^ OPC nuclei from brain tissue (*50, 59*). Control or *Atrx*^Sox10Cre^ forebrain tissue was homogenized in 500 μL homogenization buffer (20 mM Tricine KOH, 25 mM MgCl_2_, 250 mM sucrose, 1 mM DTT, 0.15 mM spermine, 0.5 mM spermidine, 0.1% IGEPAL-630, 1x protease inhibitor cocktail (Millipore Sigma Cat# 11873580001), 1 μL/mL RNase inhibitor (Thermo Fisher Scientific Cat# 10777019). Samples were diluted to 7.5 mL with homogenization buffer and filtered through a 40 μm strainer. Filtered samples were layered on top of 7.5 mL cushion buffer consisting of 0.5 mM MgCl_2_, 0.88 M sucrose, 0.5 mM DTT, 1x protease Inhibitor cocktail (Millipore Sigma Cat# 11873580001), 1 μL/mL RNase inhibitor (Thermo Fisher Scientific Cat# 10777019) and centrifuged at 2800 g for 20 mins at 4°C. Nuclei were collected as a pellet, incubated for 10 min in 500 μL 4% FBS, 0.15 mM spermine, 0.5 mM spermidine, 1x protease inhibitor cocktail (Millipore Sigma Cat# 11873580001) and 1 μL/mL RNase inhibitor (Thermo Fisher Scientific Cat# 10777019) in PBS and resuspended by gentle pipetting. Nuclei were sorted using a Sony SH800 Cell Sorter and Sun1GFP^+^ nuclei were collected. Total RNA was immediately isolated from GFP+ nuclei with a single cell RNA purification kit (NorgenBiotek Cat# 51800).

### Quantitative reverse transcriptase-PCR (qRT-PCR)

cDNA was prepared using 100 ng RNA from FANS-purified GFP^+^ OPC nuclei or from optic tracts (RNeasy Mini kit; Qiagen Cat# 74104) and 1.5 μL of 100 ng/L of random hexamers (Integrated DNA Technologies Cat# 51-01-18-26) were diluted to a final volume of 12 μL with RNAse free water. Samples were heated at 65°C for 10 min followed by addition of 4 μL first strand buffer (Thermo Fisher Scientific Cat# 18064014), 2 μL 100 mM DTT (Thermo Fisher Scientific Cat# 18064014), 0.8 μL 25 mM dNTPs, 0.5 μL RNaseOut (Thermo Fisher Scientific Cat# 10777019), 0.5 μL SuperScript II Reverse Transcriptase (Thermo Fisher Scientific Cat# 18064014) and 0.5 μL RNAse free water per sample. Samples were then incubated at 30°C for 10 minutes and 42°C for 45 minutes and stored at −20°C. cDNA was amplified with iQ SYBR Green Master Mix (BioRad Cat# 1708884) using the standard curve Ct method of quantification. Experiments were performed on a Chromo-4 thermocycler (MJ Research/BioRad) and analyzed with Opticon Monitor 3 and GeneX (BioRad) software. Technical duplicates were completed for each sample. Conditions for amplification were as follows: 40 cycles of 95°C for 10 seconds, 55-60°C for 20 seconds, 72°C for 30 seconds, and a final melting curve generated in increments of 0.5°C per plate read. Primer sequences are listed in **Supplementary Table 1**.

### RNA-seq analysis

RNA-seq libraries were prepared with 90 ng of RNA using VAHTSTM Total RNA-seq (H/M/R) Library Prep Kit for Illumina (Vazyme NR603-01) following the manufacturer’s instructions. Libraries were sequenced at Canada’s Michael Smith Genome Sciences Centre (BC Cancer Research, Vancouver, BC, Canada) using the Illumina HiseqX (Illumina Inc., San Diego, CA). On average, 50 million paired-end reads (150 bp) were obtained for each library. Raw reads were pre-processed with Trimgalore version 0.5.0, a wrapper tool around the sequence grooming tool cutadapt (*60*) with the following quality trimming and filtering parameters (‘–phred33 -length 36 -q 5 -stringency 1 -e 0.1‘). Each set of paired-end reads was mapped against the Mus musculus GRCm38.p6 primary assembly downloaded from Ensembl (*61*) release 94 (https://useast.ensembl.org/Mus_musculus/Info/Annotation) using HISAT2 version 2.0.4 (*62*). SAMtools was then used to sort and convert SAM files. The read alignments and Mus musculus GRCm38 genome annotation were provided as input into StringTie v1.3.3 (*63*) which returned gene and transcript abundances for each sample. We imported coverage and abundances for transcripts into R using the tximport (*64*) R/Bioconductor package and conducted differential analysis of transcript count data using the DESeq2 R/Bioconductor package (*65*). We used the independent hypothesis weighting (IHW) method (*66*) to weight P values and adjust for multiple testing using the procedure of Benjamini Hochberg (BH) (*67*), transcripts p-values were then aggregated using the Lancaster method (*68*). To generate the heatmap, gene level counts from the Stringtie results were transformed using variance stabilizing transformation (VST) method in DESeq2 (*65*). Z scores were then plotted using the pheatmap R package. Gene Ontology analysis was performed using Protein Analysis Through Evolutionary Relationships (Panther), http://www.pantherdb.org (*69*). Gene cluster analysis was performed using TOPPCluster (*70*).

### Chromatin immunoprecipitation (ChIP)

For ATRX ChIP, cortices from fifteen P0.5 pups were dissected and OPCs were cultured for 9 DIV in a mixed glial culture. OPCs were then physically separated from other cells by shaking at 220 rpm overnight in a tissue culture incubator at 5% CO_2_ (*57, 58*). OPCs were pooled, homogenized and passed through a 70 μm strainer (Fisher Cat#22363548). For H3K27Ac ChIP, 3 to 4 optic tracts were dissected from either control or *Atrx*^Sox10Cre^ P20 mice. Tissue was minced and triturated in DMEM and filtered through a 70μm strainer (Fisher Cat#22363548).

OPCs were first crosslinked with 0.002M ethylene glycol bis(succinimidyl succinate) (EGS, Thermo Fisher Scientific Cat # 21565) at room temperature for 45 min with constant mixing (*71*). Both OPC and optic tract samples were crosslinked with 1% formaldehyde at room temperature for 20 minutes (OPCs) or 10 minutes (optic tracts) with constant mixing. Samples were quenched in 0.125 M glycine for 2 minutes at room temperature and centrifuged at 700 g for 5 minutes. Pellets were washed twice with 10 mL cold PBS and stored at −80°C.

Chromatin was sonicated in a Bioruptor Pico Sonication Device (Diagenode Cat# B01060010) for 6 cycles (OPCs) or 13 cycles (optic tracts). For OPCs, protein G Dynabeads (50 μL, Invitrogen Cat# 10009D) were loaded with either 10 μg anti-ATRX rabbit polyclonal antibody (Abcam Cat# ab97508, RRID:AB_10680289) or anti-IgG rabbit polyclonal antibody (Thermo Fisher Scientific Cat# 02-6102, RRID:AB_2532938) for 4 hours with rotation at 4°C. For optic tracts, protein G Dynabeads (20 μL, Invitrogen Cat# 10009D) were loaded with either 2.5 μg anti-H3K27Ac rabbit polyclonal antibody (Abcam Cat# ab4729, RRID:AB_2118291) or anti-IgG rabbit polyclonal antibody (Thermo Fisher Scientific Cat# 02-6102, RRID:AB_2532938) for 1 hour with rotation at 4°C.

Chromatin lysates (50 μg per IP for OPCs, 20ug per IP for optic tracts) were precleared with 100 uL (OPCs) or 20ul (optic tracts) protein G Dynabeads (Invitrogen Cat# 10009D) for 1 hour at 4°C. Precleared lysate and antibody-bound beads were combined and incubated at 4°C overnight with rotation. The next day, beads were washed five times and eluted for 15 minutes at room temperature followed by a second elution for 20 minutes at 65°C (*71*). Crosslinking was reversed by adding 5M NaCl at 65°C for 4 hours. Samples were treated with 1 μL 10 mg/mL RNase A at 37°C for 30 min then 1 μL 20 μg/mL proteinase K for 1 hour at 45°C. 1/10^th^ of precleared chromatin lysate was treated the same and used as input.

### ChIP-seq analysis

ChIP-seq libraries were made with Accel-NGS 1S Plus DNA library preparation kit (Swift Biosciences) following the manufacturer’s instructions. Libraries were sequenced with Illumina NovaSeq 6000 and ~25 million paired-end reads (150 bp) were obtained from each library. The paired-end reads in Fastq format were trimmed with Trim Galore (v.0.6.6), a wrapper tool around Cutadapt (*60*) package, and were then aligned to the mouse genome (mm10) using Bowtie2(v.2.4.4) with the default parameters (*72*). SAMtools (v.1.12) was then used to create and sort BAM files from the aligned reads recorded in SAM format (*73*). The duplicated reads were marked with MarkDuplicates function from Picard (v.2.26.3). The mitochondrial DNA reads and blacklist regions of the genome were filtered out using Bedtools intersect (v.2.30.0). MACS2 (v.2.2.7.1) (*74*) broadPeak mode was used to call the peaks from the filtered BAM files with input control. R package csaw (v.1.26.0) (*75*) was used to count the reads in 300bp non-overlapping windows. Background noise was estimated by counting reads in 2000bp bins. We selected 300bp windows that have a signal higher than log2(3) above background. Windows were then merged if less than 100bp apart but did not extend above 5kb width. Rtracklayer (v.1.52.1) (*76*) package was then used to export the filtered windows in a BED format. FindMotifsGenome.pl and annotatePeaks.pl functions from HOMER (v.4.10) (*77*) were used to find enriched DNA motifs in the peak lists and annotate the peaks in the genome, respectively. The ATRX ChIP Bigwig track was generated using Deeptools2 bamCompare (v.3.5.2) (*78*) with the parameter “—scaleFactorsMethod SES” (input control was used for normalization). plotHeatmap and plotProfile functions of Deeptools2 were also used to compare ATRX enrichment at peaks of publicly available ChIP-seq datasets. H3K27Ac and Olig2 ChIP data was downloaded from NCBI Gene Expression Omnibus (GSE42454). UCSC liftOver tool was used to convert peak coordinates from rn4 to mm10 genome assembly. The raw fastq files of HDAC3 and p300 in OPCs were downloaded from NCBI GSE76412. The reads were aligned to rn4 genome assembly using Bowtie2 (v.2.4.4). SAMtools (v.1.12) was then used to generate and filter BAM files. Peaks were called as described for ATRX ChIP. UCSC liftOver was then used to convert peak coordinates from rn4 to mm10 genome assembly. Chd7 and Chd8 ChIP peaks in OPCs were downloaded from NCBI GSE116601. Brn1, Oct6, and Sox2 ChIP-seq data from mouse neural stem cell line NS5 was downloaded (in wig format) from NCBI GSE69859. Wig files were converted to bigwig files using wigToBigWig from rtracklayer (v.1.52.1). Bigwig files were then converted to bedGraph files using UCSC bigWigToBedGraph tool. UCSC liftOver was then used to convert peak coordinates from mm8 to mm10 genome assembly.

### Statistical analysis

Statistical analysis was performed using GraphPad Prism6 software (6.05; GraphPad Software Inc.) and all results are expressed as the mean ± SEM. Two independent data sets were compared with the Student’s t test (unpaired, 2-tailed). Multiple independent data sets were compared with a one-way ANOVA with post-hoc Tukey’s test. P values of 0.05 or less were considered to indicate significance.

## Data availability

The datasets for the *Atrx*^Foxg1Cre^ RNA microarrays were deposited in the National Centre for Biotechnology Information Gene Expression Omnibus Database (GSE210863). ATRX OPC nuclei RNA-seq and ATRX OPC ChIP-seq data can be accessed at NCBI Bioproject PRJNA866270. Other datasets used in this study are GSE3243610 (ChIP-seq OPC CHD7 and CHD8), GSE1040153 (ChIP-seq OPC Olig2, H3K27Ac), GSE1988947 (ChIP-seq OPC HDAC3), GSE1988950 (ChIP-seq OPC P300) and GSE69859 (ChIP-seq NSC, Sox2, Brn-1 and Oct6).

## Study approval

All procedures involving animals were conducted in accordance with the regulations of the Animals for Research Act of the Province of Ontario and approved by the University of Western Ontario Animal Care and Use Committee.

## Abbreviations

ATRX: α-Thalassemia/mental retardation X-linked
NSC: neural stem cell
OL: oligodendrocyte
iOL: immature oligodendrocyte
mOL: mature oligodendrocyte
OPC: oligodendrocyte precursor cell
priOPC: primitive oligodendrocyte precursor cell
Olig2: oligodendrocyte transcription factor 2
PDGFRα: platelet derived growth factor receptor α
HDAC3: histone deacetylase 3
CNS: central nervous system
ID: intellectual disability
Swi/Snf: Switch/Sucrose non-fermenting
Nlgn4: Neuroligin 4
T4: thyroxine
T3: triiodothyronine
GFP: green fluorescent protein
P: postnatal day
FANS: fluorescent activated nuclei sorting
qRT-PCR: Quantitative reverse transcriptase-PCR
co-IP: co-immunoprecipitation
ChIP: chromatin immunoprecipitation
MAG: myelin associated glycoprotein
MOG: myelin oligodendrocyte glycoprotein
MBP: myelin basic protein
PLP1: proteolipid protein 1
PBS: phosphate buffered saline
ANOVA: analysis of variance
APC: adenomatous polyposis coli
IP: intraperitoneal
DIV: days in vitro
NG2: neural glial antigen 2
S100β: S100 calcium-binding protein β
GFAP: glial fibrillary acidic protein
Ctx: cortex
CC: corpus callosum
Ctrl: control
SC: subcutaneous
Tam: tamoxifen
IGF-1: insulin-like growth factor 1.

## Results

### Decreased myelination in the Atrx^FoxG1Cre^ mouse forebrain

We previously reported that postnatal health and longevity are adversely affected in mice with ATRX deficiency in the forebrain, anterior pituitary and liver (*Atrx*^FoxG1cre^ mice) (*55, 79*). These mice have a reduced lifespan, growth and hormone abnormalities, including low circulating levels of IGF-1 and T4 (*55, 79*) and exhibit aberrant gene expression in the liver (*55*). To identify potential forebrain related differences in transcription between control and *Atrx*^FoxG1Cre^ mice, we performed a microarray analysis of 3 littermate-matched pairs of P17 control and *Atrx*^FoxG1Cre^ mice. Microarray analysis revealed a significant downregulation of genes involved in myelination and oligodendrocyte morphology (**Figure 1a,b**, p<0.05). Moreover, downregulated genes are highly expressed in mouse oligodendrocytes according to the Allan Brain Atlas single cell RNA gene sets (**Figure 1c**). Given previous reports of white matter abnormalities in ATR-X syndrome patients (*33, 34*), we further explored whether the mice exhibit a similar phenotype. We examined the level of well-established myelin markers by immunofluorescence staining of brain cryosections. The results showed decreased level of myelin oligodendrocyte glycoprotein (MOG) and myelin basic protein (MBP) in the corpus callosum and adjacent cortex in *Atrx*^FoxG1Cre^ compared to control mice at P20 (**Figure 1d**). Western blot analysis of myelin associated glycoprotein (MAG), MOG and MBP showed that myelin protein expression is significantly decreased in *Atrx*^FoxG1Cre^ forebrain compared to littermate matched controls (n=4 each genotype; Student’s T-test, MAG p=0.006, MOG p=0.005, MBP p=0.009) (**Figure 1e,f)**. These data strongly suggest that the *Atrx*^FoxG1Cre^ CNS is hypomyelinated.

**Figure 1.**
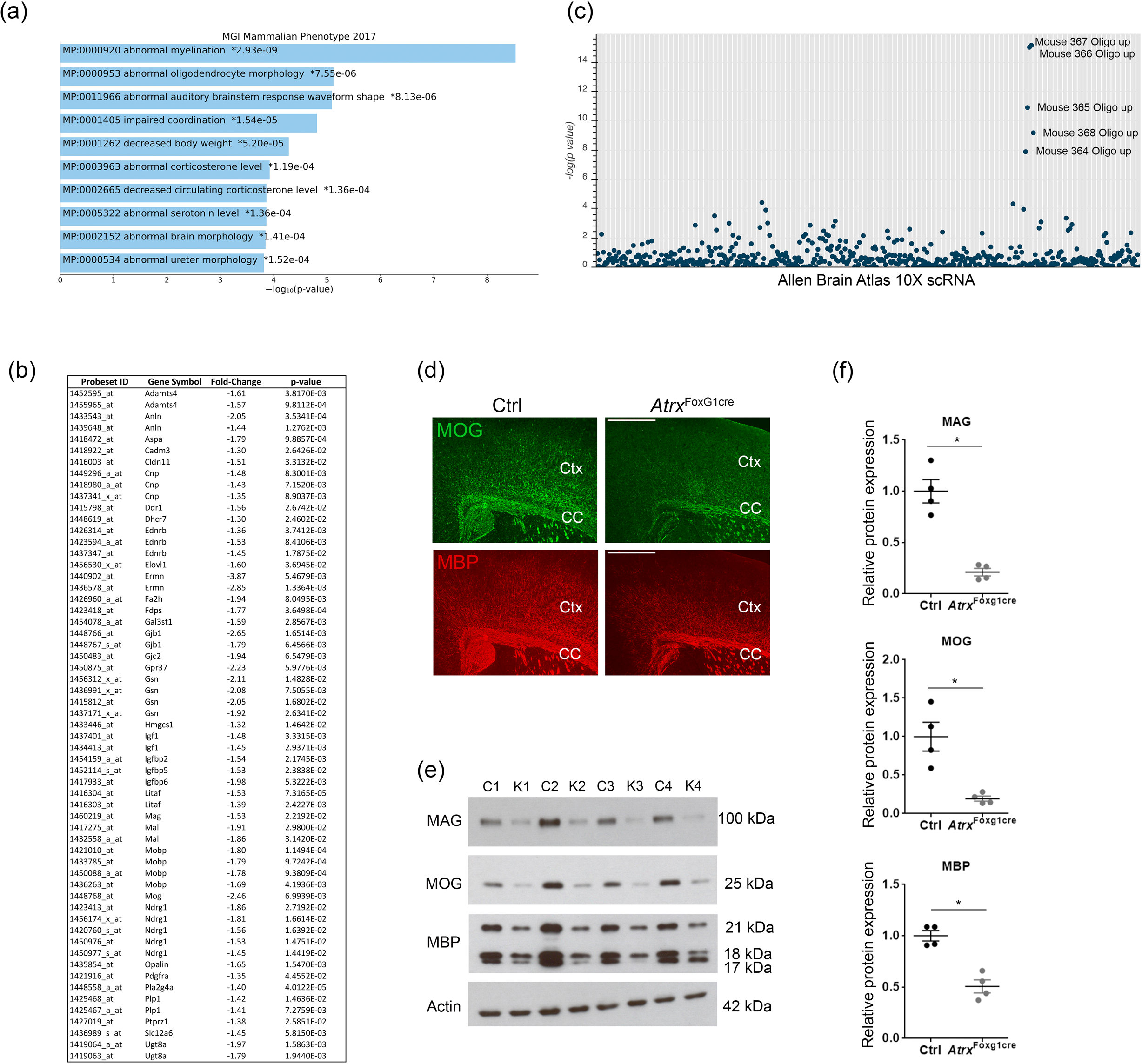
Defective myelination in the ATRX-null mouse forebrain. (a) Microarray analysis of P17 control and *Atrx*^FoxG1cre^ forebrain tissue reveals a significant downregulation (P<0.05) of genes related to abnormal myelination and oligodendrocyte morphology. (b) List of genes corresponding to the “Abnormal Myelination” category, along with fold change and p-value (c) Manhattan plot of Allan Brain Atlas 1 OX scRNA gene set enrichment from the list of downregulated genes. Each line on the x-axis denotes a single gene set and the y-axis measures the −log,_10_(p-value) for each gene set. (d) lmmunofluorescence microscopy of P20 brain cryosections stained with anti-MOG (green) and anti-MBP (red) antibodies confirms decreased levels of these myelin proteins in the cortex (Ctx) and corpus callosum (CC) areas of the forebrain in *Atrx*^FoxG1cre^ mice compared to controls (Ctrl). Scale bar, 500 μm. (e,f) Western blot analysis of P20 forebrain protein extracts showing significantly decreased levels of MAG, MOG and MBP in *Atrx*^FoxG1cre^ compared to control mice (n=4 for each genotype) after normalization to β-actin protein levels. Error bars represent+/− SEM and asterisks indicate p<0.05 (Student’s t-test).

### Postnatal thyroxine treatment partially restores myelination deficits in Atrx^FoxG1Cre^mice

Circulating levels of the thyroid hormone T4 in *Atrx*^FoxG1Cre^ mice are less than half that of controls, most likely stemming from the deletion of *Atrx* in the anterior pituitary in these mice (*55, 79*). The thyroid hormone receptor is a transcription factor that requires its ligand, triiodothyronine (T3), to promote the expression of several myelin genes (*23–25*). We therefore tested whether postnatal administration of the prohormone T4 might rescue hypomyelination in *Atrx*^FoxG1Cre^ mice. We performed daily injections of 0.1 mg/kg L-thyroxine (T4) or PBS from P0 to P14, as outlined in **Figure 2a**, a dose previously shown to successfully elevate circulating T4 to control levels in *Atrx*^FoxG1Cre^ mice (*55*).

**Figure 2:**
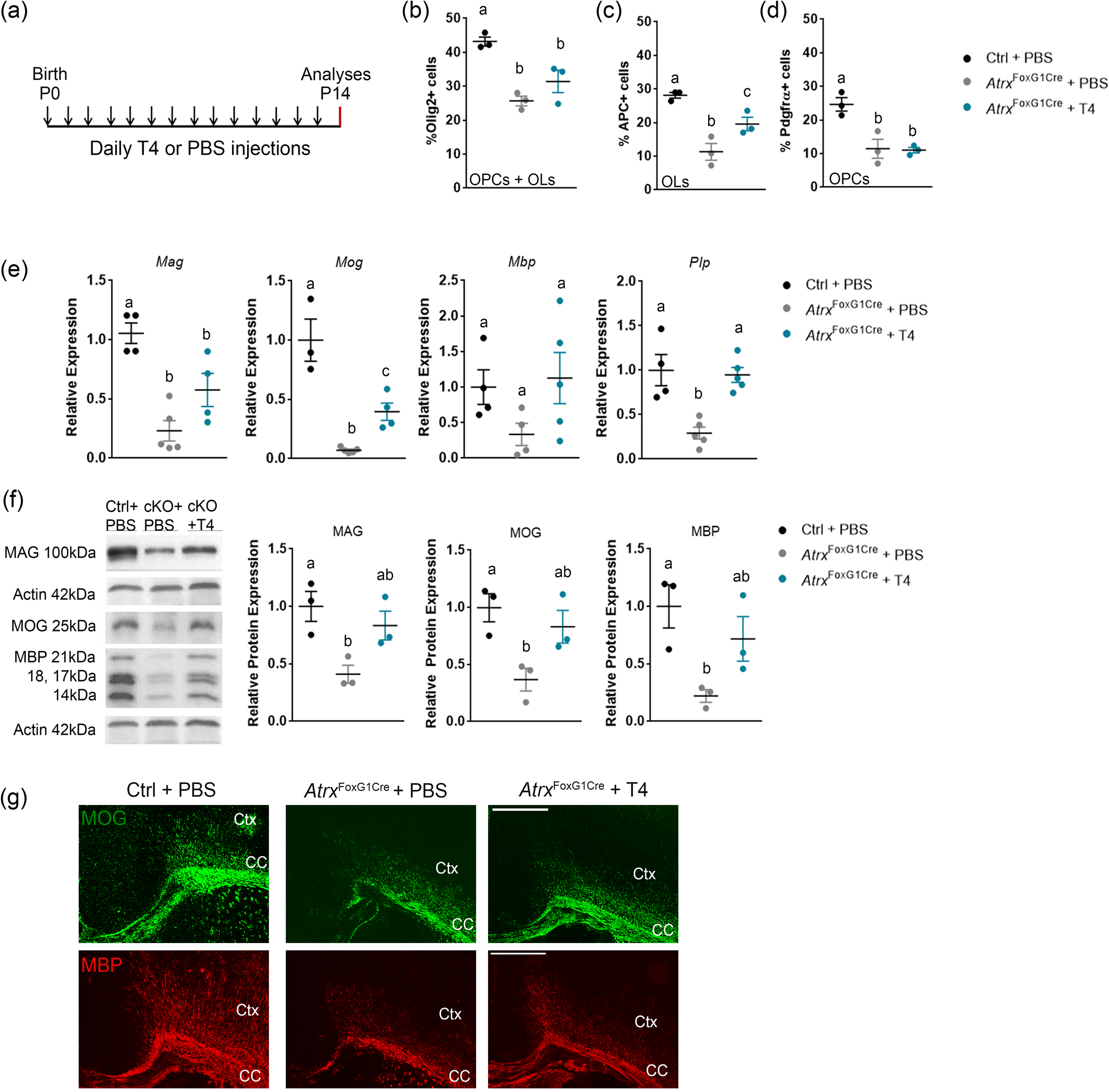
Postnatal thyroxine treatment partially rescues the number of APC^+^ Ols and the expression of myelin proteins in the ATRX-null forebrain but does not restore the number of Pdgfraα^+^ OPCs. (a) Outline of experimental design. Each arrow represents a subcutaneous (SC) injection of 0.1 mg/kg T4 or PBS to *Atrx*^FoxG1cre^ and control pups. Analyses were then performed at P14. (b-c) Quantification of immunostained P14 cryosections demonstrate that the reduced number of Olig2α^+^ and Pdgfraα^+^ cells observed in the *Atrx*^FoxG1cre^ CC is not rescued by T4 treatment (n=3 each group). (d) The number of APCα cells are partially rescued upon T4 treatment (n=3 each group). (e) qRT-PCR shows that transcript levels of several myelin genes are decreased in the P14 AtrxFoxG1cre forebrain and partially restored to control levels following T4 treatment (n=3-5 for each group). (f) Western blot analysis of MAG, MOG and MBP shows decreased protein levels in P14 Atrx^FoxGlcre^ compared to control forebrain extracts and partial rescue upon T4 treatment. Graphs show quantification of 3 trios, normalized to Actin protein levels. (g) lmmunofluorescence staining of P14 brain cryosections highlight the decreased expression of MOG (green) and MBP (red) protein levels in the *Atrx*^FoxG1cre^ cortex (Ctx) and corpus callosum (CC) that are ameliorated following T4 treatment. Scale bar, 500 μm. In all graphs, error bars represent +/−SEM, and groups with different letters significantly differ from one another (one-way ANOVA with post-hoc Tukey HSD; p<0.05).

We first quantified the number of OLs after T4 treatment by immunofluorescence staining of P14 brain cryosections, using an antibody specific for the pan-OL marker Olig2. We observed a significant reduction of the proportion of Olig2^+^ cells (representing all OPCs and OLs) in the *Atrx*^FoxG1Cre^ corpus callosum (25.7% of total DAPI^+^ cells) compared to controls (43.2% of total DAPI^+^ cells) and that this was partially recovered in *Atrx*^FoxG1Cre^ mice treated with T4 (31.4%) (n=3 for each genotype; p=0.004, one-way ANOVA) (**Figure 2b**). We also counted the proportion of OPCs or OLs in the corpus callosum of control and *Atrx*^FoxG1Cre^ brain cryosections at P14, marked by PDGFRα and adenomatous polyposis coli (APC), respectively. The proportion of differentiated APC^+^ OLs was decreased in the *Atrx*^FoxG1Cre^ corpus callosum (11.3%) compared to controls (28.2%) and was partially recovered (19.6%) following T4 treatment (n=3 for each genotype; p=0.002 one-way ANOVA) (**Figure 2c**). While the proportion of PDGFRα^+^ OPCs is lower in *Atrx*^FoxG1Cre^ mice (11.5%) compared to controls (24.6%), it was not improved upon T4 treatment (11%) (n=3 each genotype; p=0.005, one-way ANOVA) (**Figure 2d**), suggesting that thyroxine can only remediate differentiation defects.

To confirm that T4 administration promotes the differentiation of ATRX-null OPCs, we performed qRT-PCR for several established myelin genes including *Mag, Mog, Mbp* and *Plp1* (*Proteolipid protein 1*). The level of these RNA transcripts was elevated in T4-vs PBS-treated *Atrx*^FoxG1Cre^ mice, but the extent of rescue did not reach control levels for *Mag* and *Mog* (n=3-4 each genotype; *Mag* p=0.001; *Mog* p=0.0001, *Mbp* p=0.17, *Plp* p=0.001 by one-way ANOVA) (**Figure 2e**). Similarly, Western blot analysis of MAG, MOG and MBP showed some rescue of myelin protein levels in T4-treated *Atrx*^FoxG1Cre^ forebrain extracts (n=3 each genotype MAG p=0.026; MOG p=0.027, MBP p=0.035 by one-way ANOVA) (**Figure 2f**). Immunofluorescence staining of MOG and MBP in *Atrx*^FoxG1Cre^ forebrain cryosections also showed improved myelination after T4 administration (**Figure 2g**). Taken together, these results suggest that hypomyelination in *Atrx*^FoxG1Cre^ mice is due to a reduced number of OPCs and OLs and that T4 treatment stimulates OPC differentiation without increasing OPC numbers, which could explain the incomplete effect of T4 treatment on the extent myelination.

### Targeted ablation of Atrx in neurons does not affect myelination

The results of the T4 experiments suggested that ATRX is required in the central nervous system to support normal myelination, but it was not clear whether this effect is cell autonomous in OPCs or caused by ATRX loss in other cell types that provide trophic support to OPCs. To resolve whether ATRX regulates myelination intrinsically in OPCs or indirectly via neuronal signaling (or both), we generated mice that lack ATRX in forebrain excitatory neurons using the *NexCre* driver mice for *Atrx*^loxP^ recombination (*48*). We confirmed the absence of ATRX nuclear protein in cortical neurons by co-immunofluorescence staining of ATRX and the neuronal marker NeuN (**Figure S1a**). We next examined myelin protein expression in the forebrain by immunofluorescence staining of P20 brain cryosections and no obvious difference in MBP and MOG protein was observed between control and *Atrx*^NexCre^ mice (**Figure S1b**). Furthermore, MAG, MOG and MBP protein levels were equivalent to controls when assessed by immunoblot of P20 *Atrx*^NexCre^ forebrain extracts (n=3 each genotype; Student’s T-test, MAG p=0.627; MOG p=0.917, MBP p=0.425) (**Figure S1c**). These results establish that ATRX is not required in neurons of the developing brain for proper signaling to OPCs and for normal production of myelin.

### Loss of ATRX in OPCs leads to hypomyelination

We first verified that ATRX is expressed in OPCs and OLs in the mouse brain by co-staining of ATRX with PDGFRα and APC, respectively (**Figure S2**). Next, we targeted ATRX deletion in primitive OPCs (priOPCs) and OPCs using the tamoxifen-inducible *Sox10CreER^T2^* mice, which upon mating with *Atrx*^LoxP^ mice produce *Atrx*^Sox10-iCreERT2^ (*Atrx*^Sox10Cre^) mice (*49*). The Cre-sensitive nuclear membrane *Sun1GFP* reporter was also introduced to allow fate tracking of Cre-expressing cells (**Figure 3a**) (*50*). Cre recombination was induced by daily intraperitoneal (IP) injection of nursing dams for three successive days (pups at P1-P3) (*56*) and the mice were analyzed at P20. *Atrx* depletion was confirmed by isolating nuclei from control and *Atrx*^Sox10Cre^ P20 forebrains and sorting the Sun1GFP^+^ OPC nuclei. qRT-PCR analysis confirmed a 74.2% decrease in *Atrx* transcript levels in *Atrx*^Sox10Cre^ Sun1GFP^+^ nuclei compared to control Sun1GFP^+^ nuclei (n=4 each genotype; Student’s T-test, p=0.009) (**Figure 3a**).

**Figure 3:**
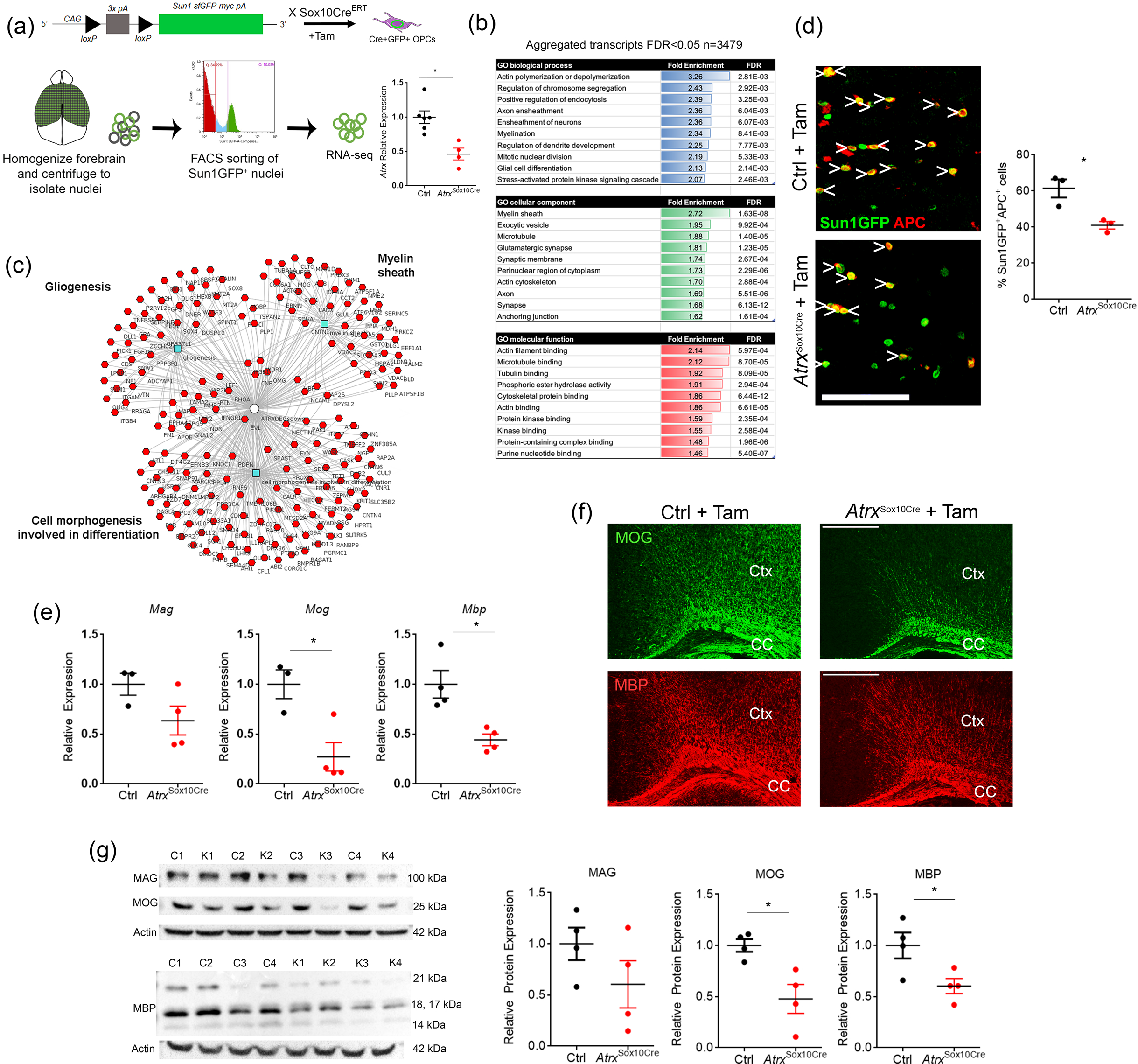
Defective differentiation of ATRX-null OPCs causes hypomyelination. (a) Sun1GFP+ nuclei were sorted and used for RNA-sequencing. (b) Pathway analysis (c) Downregulated genes (d) Cell counts show significantly fewer Sun1GFP^+^ cells co-staining with the mature OL marker APC (red) in the P20 *Atrx*^sox10cre^ corpus callosum compared to controls. Scale bar, 100 μm (n=3 each genotype; p<0.05, Student’s T-test). (e) RT-qPCR analysis shows reduced expression of Mag, Mog and Mbp in *Atrx*^sox10cre^ P35 optic tract compared to controls (n=3-4 each genotype; p<0.05, Student’s T-test). (f) lmmunofluorescence staining reveals that myelin proteins MOG (green) and MBP (red) are reduced in P20 *Atrx*^sox10cre^ cortex (Ctx) and corpus callosum (CC) compared to controls. Scale bar, 500 μm. (g) Western blot analysis of MAG, MOG and MBP (left) and quantification (right) confirms decreased expression of these proteins in P20 *Atrx*^sox10cre^ mouse forebrain extracts compared to controls after normalization to β-actin (n=4 each genotype; p<0.05 Student’s T-test). In all graphs, error bars represent +/−SEM and asterisks indicate significance of p<0.05.

We next performed RNA-seq of Sun1GFP+ nuclei obtained from control and *Atrx*^Sox10Cre^ forebrain and identified 3479 altered genes (aggregated transcripts, Lancaster method, p value<0.05). Gene ontology analysis identified categories of enrichment for biological processes “myelination”, “axon ensheathment” and “ensheathment of neurons”, as well as “glial cell differentiation” and “actin polymerization and depolymerization”. Moreover, the top cellular component category of enrichment was “myelin sheath” and many of the molecular function enrichment categories featured aspects of cytoskeletal biology, possibly reflecting extensive changes in cell shape and membrane architecture during myelination (**Figure 3b**). Cluster analysis of only the downregulated genes further emphasized that numerous genes implicated in myelination, gliogenesis and cell morphogenesis involved in differentiation are downregulated in ATRX-null priOPCs and their progeny (**Figure 3c**).

To verify potential defects in OPC differentiation and myelination, we performed immunofluorescence staining for APC, a marker of differentiated but immature OLs (iOLs) in P20 control and *Atrx*^Sox10Cre^ cortical sections. There was a significant reduction in the number of Sun1GFP^+^ ATRX-null cells that co-expressed APC (41%) compared to Sun1GFP^+^ control cells (61.4%) (**Figure 3d**) (n=3 each genotype; p=0.020 Student’s T-test), indicating a lower number of differentiating iOLs. RNA transcripts of mature OLs (mOL)-specific genes *Mag, Mog, Mbp* were also significantly decreased in P35 *Atrx*^Sox10Cre^ optic tract (n=3-4 each genotype; *Mag* p=0.119 *Mog* p=0.017, *Mbp* p=0.010 Student’s T-test) (**Figure 3e**). Furthermore, immunofluorescence staining for MOG and MBP in brain cryosections at P20 showed decreased level of these myelin proteins in the corpus callosum and cortex of *Atrx*^Sox10Cre^ mice compared to controls (**Figure 3f**). Reduced myelin protein expression in *Atrx*^Sox10Cre^ mice was confirmed by western blot analysis of MAG, MOG and MBP (n=4 each genotype; MAG p=0.208, MOG p=0.015, MBP p=0.036 Student’s T-test) (**Figure 3g**). These results demonstrate that ATRX-null priOPCs/OPCs have a reduced capacity to differentiate into iOLs and mOLs, leading to reduced myelination.

### Loss of the Olig2 fate specification factor in ATRX-null cells

At all stages of development, priOPCs, OPCs and OLs express Olig2, an essential regulator of OL lineage specification (*3, 80, 81*). Olig2 suppresses astrocytic fate (*5–7*) and promotes OL differentiation via regulation of the chromatin remodelling complex Smarca4/Brg1 (*10*). The absence of ATRX protein was confirmed in *Atrx*^Sox10Cre^ OPCs by immunofluorescence staining and imaging in parallel with the Sun1GFP reporter (**Figure 4a**). Upon staining the *Atrx*^Sox10Cre^ corpus callosum with Olig2 to confirm specific deletion in the oligodendrocyte lineage cells, we were surprised to see that a large proportion of Sun1GFP^+^ ATRX-null cells did not express Olig2 (47.9%) compared to Sun1GFP^+^ control OPCs (12.1%) (n=3-4 each genotype; Student’s T-test, p=0.0004, **Figure 4b,c**). Similarly, qRT-PCR of RNA isolated from sorted *Atrx*^Sox10Cre^ Sun1GFP^+^ nuclei confirmed a 58% reduction in *Olig2* transcripts compared to Sun1GFP^+^ control nuclei (n=3 each genotype; Student’s T-test, p=0.015) (**Figure 4d**).

**Figure 4:**
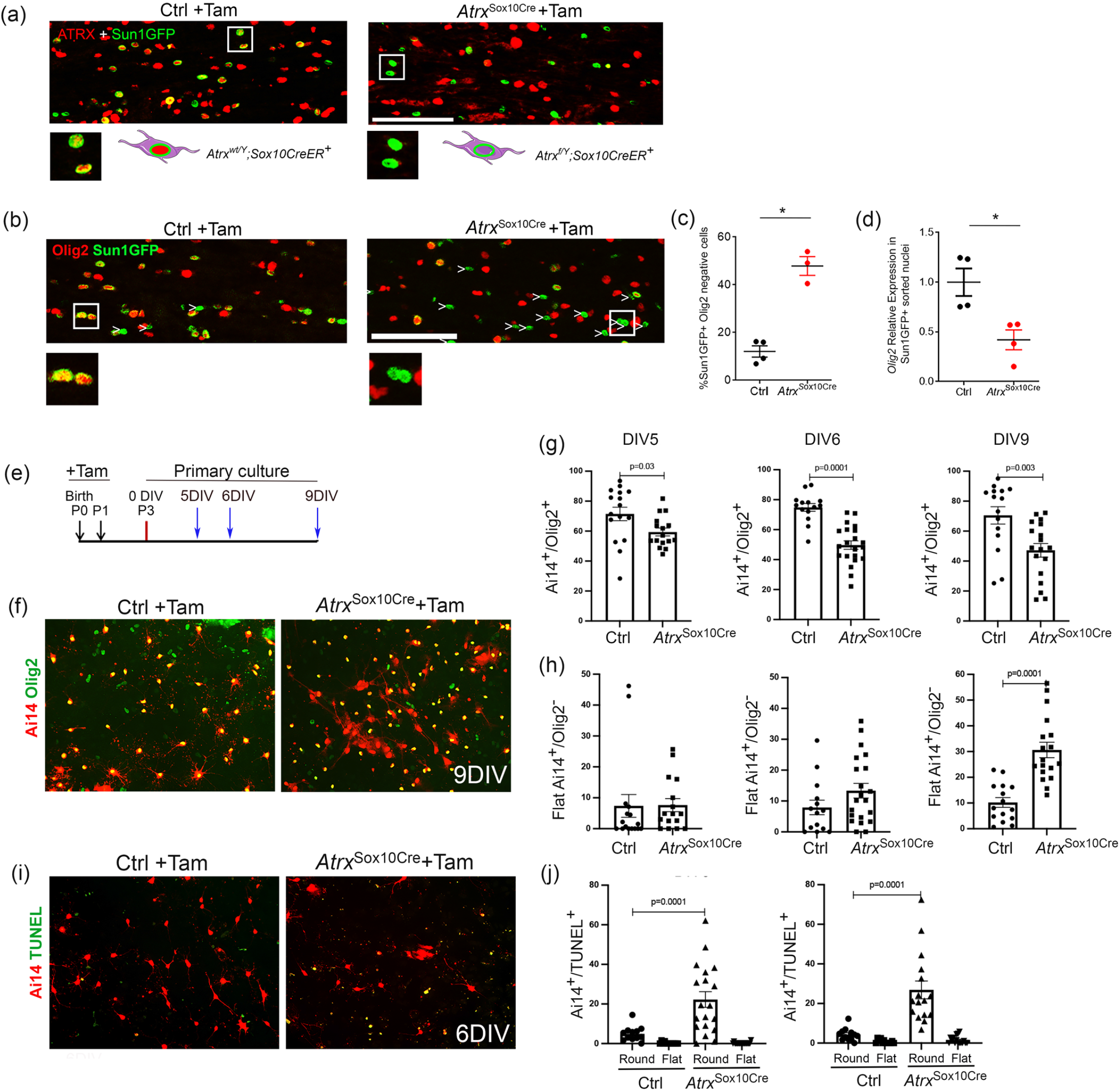
Loss of Olig2 expression and morphological changes in a subset of ATRX-null OPCs. (a) lmmunofluorescence microscopy of P20 Atrx^wt/y^;Sox10Cre^+^ corpus callosum shows expression of the Sun1GFP fusion protein in the nuclear membrane of OPCs upon Tamoxifen treatment (left). P20 Atrx^fl/Y^;Sox10Cre^+^ treated with Tamoxifen also express Sun1GFP but lack expression of nuclear ATRX (right). Scale bar, 100 μm. (b) Fate mapping of ATRX-null Sun1GFP^+^ OPCs reveals frequent loss of the Olig2 transcription factor (white arrowheads). Scale bar, 100μm. (c) Quantification of Sun1GFP^+^ OPCs that lack Olig2 staining (right) (Ctrl n=4; Atrx^sox10cre^ n=3; Student’s t-test; p=0.004). (d) qRT-PCR analysis demonstrates significantly decreased *Olig2* transcripts in Sun1GFP+ Atrx^sox10cre^ sorted nuclei compared to Sun1GFP+ control nuclei (n= 4 for each genotype; Student’s T-test; p=0.015). (e) Experimental timeline of Tamoxifen (Tam) injections to nursing mothers at PO and P1 (black arrows). Primary mixed glial cultures of cortices were established at P3 and maintained for 9 days *in vitro* (DIV). (f) Images of Ai14 control and Atrx^sox10cre^ mixed glial culture at 9DIV stained for Olig2 (green). (g) The number of Ai14/Olig2^+^ cells is decreased in ATRX-null OPCs at DIV5, 6 and 9. (h) The number of Ai14+ ATRX-null OPCs exhibiting a flattened morphology increases over time in culture, reaching significance at DIV9. (i) Images of Ai14 control and Atrx^sox10cre^ mixed glial culture at 6DIV stained for TUN EL (green). (j) Quantification of TUNEL staining shows increased cell death at DIV5 and DIV6 in round-shaped ATRX-null OPCs.

Loss of *Olig2* expression in ATRX-null cells was also observed in primary glial cell cultures. Cre-mediated recombination was induced by administration of tamoxifen to nursing dams for two days (age of pups P0-P1). P3 cortices of *Atrx*^Sox10Cre^ or control mice that also express the Ai14-TdTomato reporter (*51*) were dissociated and cells grown in mixed glial co-culture in which OPCs are expanded on top of an astrocyte monolayer (**Figure 4e**) (*57, 58*). Immunofluorescence staining revealed that many ATRX-null Ai14^+^ cells do not express Olig2 after 5, 6 and 9 days *in vitro* (DIV) (**Figure 4 f,g**) and that these cells appear elongated and flattened instead of the typical rounded shape of OPCs (**Figure 4f,h**). We also detected a significant increase in apoptosis of Ai14+ ATRX-null rounded OPCs compared to control cells at both DIV5 and DIV6 (**Figure 4i,j**). Ki67 staining showed that the same proportion of rounded Ai14+ cells proliferate in the presence or absence of ATRX at DIV5. However, a significant decrease in Ai14+/Ki16+ cells was detected at DIV6 (**Figure S3**). Overall, these results demonstrate that a subset of ATRX-null OPCs lack Olig2 expression and either undergo apoptosis or change morphology. In addition, the proliferation of ATRX-null OPCs was not initially affected but was decreased by 6 DIV compared to control cells.

### ATRX co-occupies the Olig2 locus with other OPC fate regulators and promotes an active chromatin state

We next performed ATRX chromatin immunoprecipitation on cultured primary mouse OPCs and identified 42,820 ATRX peaks using MACS2 (*74*), with the bulk of ATRX chromatin occupancy identified two highly represented DNA motifs corresponding to Oct6 (*Pou3f1*) and Brn1 (*Pou3f3*) (**Figure 5b**) from ChIP-seq data previously done in NPCs (*82*). However, when we compared ATRX enrichment +/−3kb from Oct6 and Brn1 binding site midpoints, we only observed weak ATRX enrichment flanking these transcription factor binding sites. Interestingly, ATRX binding was more noticeable at Sox2 peaks, which was analyzed in parallel, as a control (**Figure 5c**). These results suggest that ATRX may be cooperating with these transcription factors during OPC differentiation, but to a limited extent.

**Figure 5:**
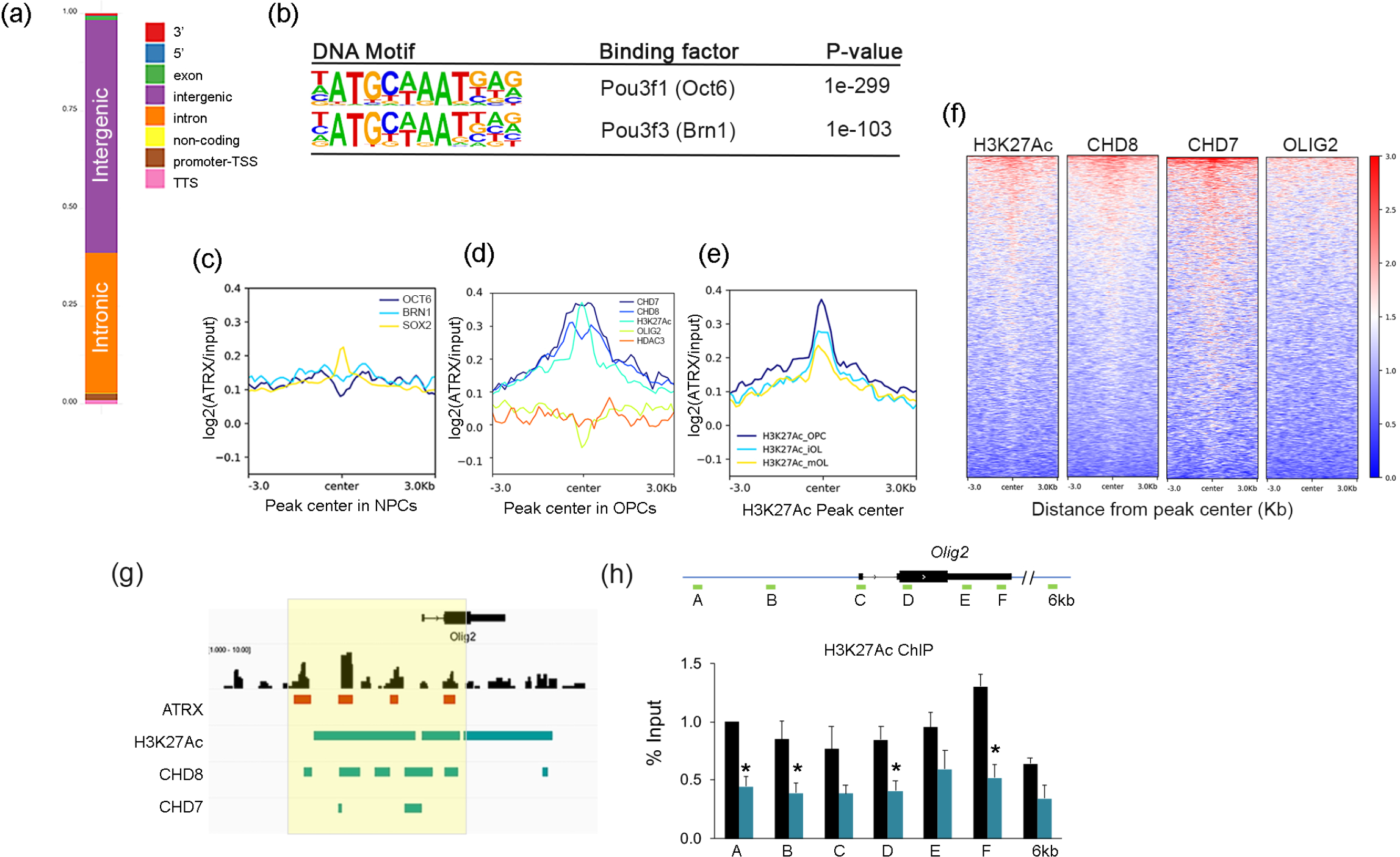
ATRX occupancy across the *Olig2* locus promotes histone H3K27Ac post-translational modification. (a) ATRX ChIP-seq in mouse primary OPCs reveals highest ATRX occupancy at intergenic sites and introns. (b)DNA motif enrichment of sequence corresponding to ATRX binding corresponds to Oct6 and Brn1 transcription factor binding sites identified in NPCs (c) ATRX genome wide occupancy at ChIP-seq peaks of Sox2, Brn1 and Oct6 in NPCs. (d) ATRX occupancy at ChIP-seq peaks of HDAC3, Olig2, CHD7, CHD8 and H3K27Ac in OPCs. (e) Co-occurance of H3K27Ac and ATRX in OPCs, iOLs and mOLs. (f) Heatmaps of ATRX occupancy at ChlP-seq peaks of H3K27Ac, CHD8, CHD7 and OLIG2 in OPCs. (g) IGV view of the *Olig2* gene showing ATRX ChIP-seq peaks (red) and how they align with H3K27Ac, CHD8 and CHD7 (green). (h) ChlP-qPCR of H3K27Ac demonstrates reduced enrichment at several sites at the *Olig2* gene in the absence of ATRX.

We then turned to other established critical factors in OPC differentiation, including HDAC3 (*83*), CHD7 (*84*), CHD8 (*85, 86*) and OLIG2 (*3, 81, 87*). Comparing ChIP-seq of ATRX and these factors in OPCs showed substantial co-occupancy of ATRX with chromatin remodelers CHD7 and CHD8, but not with HDAC3 or OLIG2 (**Figure 5d,f**). CHD7 and CHD8 are ATP-dependent chromatin remodeling proteins involved in OL differentiation and myelination (*84, 85*). CHD8 has been implicated in the establishment of OPC identity, while both CHD7 and CHD8 target enhancers of key myelinogenic genes (*84*), which are marked by histone H3 acetylated at lysine 27 (H3K27Ac) (*88*). Comparison of H3K27Ac ChIP-seq peaks from OPCs, immature OLs (iOLs) and mature OLs (mOLs) revealed elevated ATRX binding at active enhancers in these cell types, although it was more pronounced in OPCs compared to iOLs and mOLs (**Figure 5e,f**). ATRX directly binds the *Olig2* gene locus at regulatory sites defined by H3K27Ac and CHD8, but not for CHD7 (**Figure 5g**). An overlap of ChIP-seq peaks between ATRX, H3K27Ac and CHD8 is also observed at numerous other genes involved in OL fate specification and differentiation, such as *Nkx2-2*, *Sox8*, *Thra* and *Mbp* (**Figure S4**). Given the strong association between H3K27Ac and ATRX, we performed ChIP-qPCR of H3K27Ac in control and *Atrx*^Sox10Cre^ optic nerves. The results demonstrate a significant reduction of this histone posttranslational modification at the *Olig2* gene locus in the absence of ATRX, corresponding to reduced gene expression (**Figure 5h**).

We further compared the list of differentially expressed genes identified by RNA-seq, to cell type-specific gene lists (*89*) and determined that the majority of downregulated genes are highly expressed not only in OLs, but also OPCs, suggesting that perhaps OPC determination is compromised in the absence of ATRX (**Figure 6a**). In addition to *Olig2*, several other transcription factors involved in OPC fate were downregulated according to RNA-seq data, including *Olig1*, *Sox8*, and *Zfp36l1*. The latter codes for a transcription factor that promotes differentiation of multipotent progenitors into pri-OPCs (*90*) suggesting that ATRX-null cells fail to express early determinants of both OPC and pri-OPC identify (**Figure 6b**). Indeed, RT-qPCR analysis shows a reduction in the expression of the OPC markers *Pdgfrα*, *Cspg4* (NG2) and *Gpr17* (n=3-4 each genotype; Student’s T-test, *Pdgfrα* p=0.017, *Cspg4* p=0.032, *Gpr17* p=0.0004) in the *Atrx*^Sox10Cre^ mice as compared to controls. The priOPC-expressed genes *Olig1* and *Sox10* were similarly reduced but did not reach the significance threshold of p<0.05 (**Figure 6c**) (n=3 each genotype; *Olig1* p=0.081, *Sox10* p=0.133 Student’s T-test). Moreover, Sun1GFP+ ATRX-null OPCs in the corpus callosum stain less frequently for PDGFRα (24.9%) compared to controls (41.1%) (**Figure 6d,e**) (n=3 each genotype; p=0.001, Student’s T-test). Immunofluorescence staining was also performed on mixed glial cultures at different timepoints to examine OPC development over time. We observed that a significantly decreased proportion of ATRX-null Ai14+ cells expressed PDGFRα at 5, 6 and 9 DIV compared to control cells, the difference becoming more pronounced at 6 and 9DIV (**Figure 6f,g**). We conclude that ATRX ablation in postnatal Sox10-expressing priOPCs and OPCs leads to a loss of progenitor identity.

**Figure 6:**
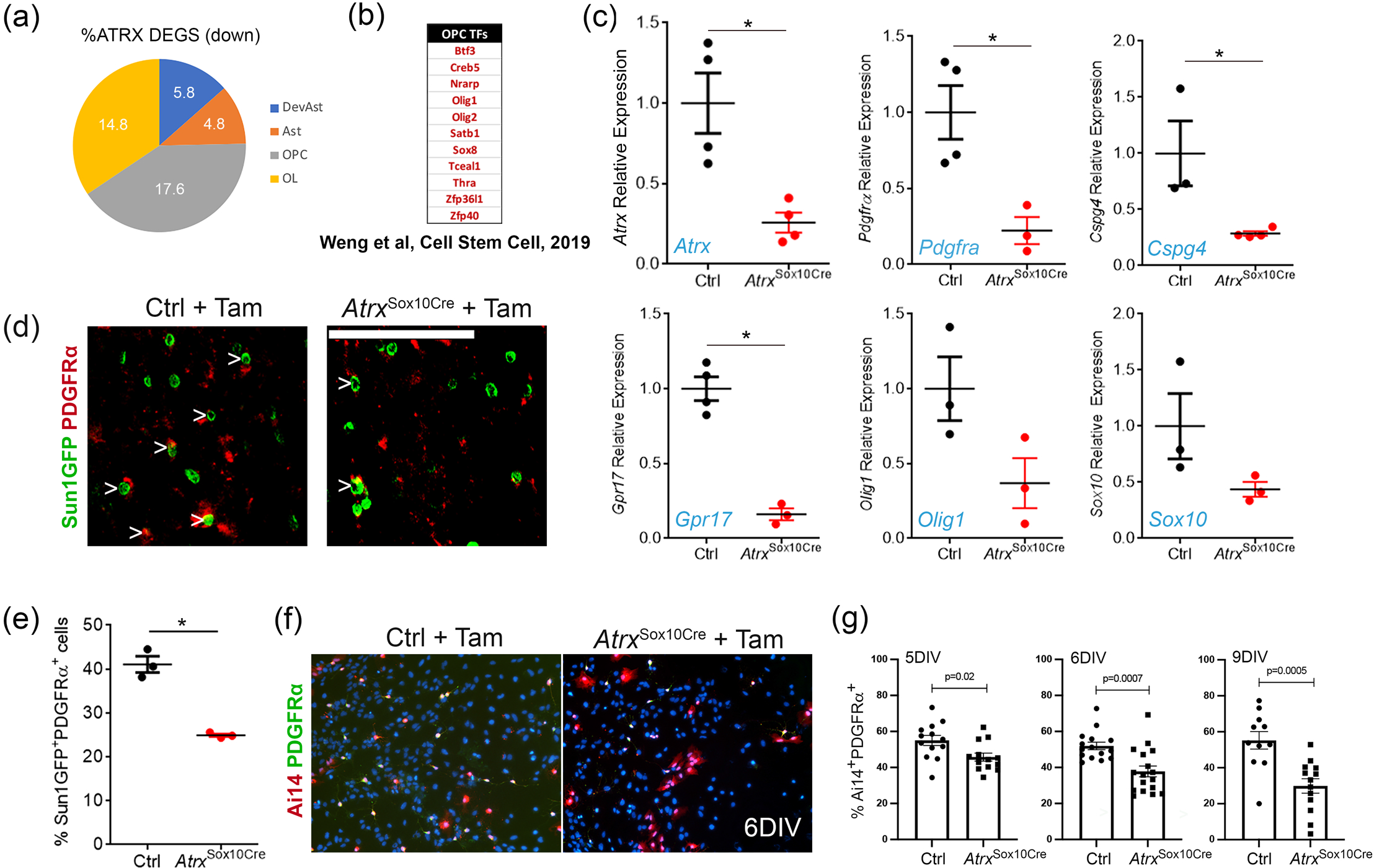
Loss of OPC identity upon *Atrx* deletion in Sox10^+^ cells *in vivo* and *in vitro*. (a) Transcriptional analysis of *Atrx*^sox10cre^ sorted Sun1GFP^+^ nuclei shows that downregulated genes are equally enriched in OLs and OPCs based on data reported by Cahoy et al, 2008. (b) Several transcription factors that icontribute to OPC fate are downregulated upon loss of ATRX, whereas astrocyte transcription factors are upregulated. (c) RT-qPCR analysis of Sun1GFP+ sorted nuclei demonstrates that key genes that specify OPCs are downregulated in the absence of ATRX (n=3-4 each genotype; asterisks indicate p<0.05, Student’s t-test). (d) PDGFRα immunostaining (red) of Sun1GFP^+^ PDGFRα^+^ cells in the corpus callosum of *Atrx*^sox10cre^ compared to control mice. Scale bar, 100 μm. (e) Quantification of Sun1GFP^+^ PDGFRα^+^ cells (n=3 each genotype; Student’s t-test; p=0.001). (f) lmmunofluorescence staining of PDGFRα (green) shows a majority of Ai14^+^ (red) cells are PDGFRα^+^ after 6DIV. (g) The percentage of Ai14+ cells that express PDGFRα was quantified after 5, 6 and 9DIV in tamoxifen-treated control (Ctrl) and *Atrx*^sox10cre^ cell culture. In all graphs, error bars represent +/−SEM.

### Loss of ATRX leads to ectopic astrogliogenesis

Ablation of the transcription factor Olig2 in OPCs in the postnatal cortex has been reported to induce differentiation into astrocytes (*5–7*). This observation, combined with the observation that many ATRX-null OPCs take on an atypical morphology reminiscent of astrocytes *in vivo* and *in vitro*, prompted us to investigate whether ATRX-null OPCs adopt a more malleable state conducive to alternative fate choice. To answer this question, we compared RNA-seq data to the list of the most enriched genes in astrocytes, and found that many astrocyte-enriched genes were upregulated in ATRX-null cells (**Figure 7a**). We then performed immunofluorescence staining of known astrocyte markers in control and *Atrx*^Sox10Cre^ brain cryosections at P20. We observed increased S100 calcium-binding protein β (S100β) staining in the corpus callosum and cortex in *Atrx*^Sox10Cre^ mice (**Figure 7b**). Cells co-expressing the endogenous Sun1GFP and S100β (by immunostaining) were counted and the results show a 20% increase in the number of Sun1GFP^+^ cells that co-express S100β in *Atrx*^Sox10Cre^ compared to control corpus callosum (n=3 each genotype; Student’s T-test, p=0.003) (**Figure 7c**). S100β can sometimes stain OLs (*91*) and indeed we observed that a proportion of control Sun1GFP^+^ OPCs express S100β (34.2%). We thus used another well-known astrocyte marker, glial fibrillary acidic protein (GFAP). Similar to the results obtained for S100β, we observed increased GFAP staining in the cortex of *Atrx*^Sox10Cre^ mice (**Figure 7d**). Cell counts of GFAP^+^Sun1GFP^+^ cells showed a 15.2% increase in the proportion of Sun1GFP^+^ cells that co-stain with GFAP in *Atrx*^Sox10Cre^ compared to control in the corpus callosum (n=3 each genotype; Student’s T-test, p=0.007) (**Figure 7e**).

**Figure 7:**
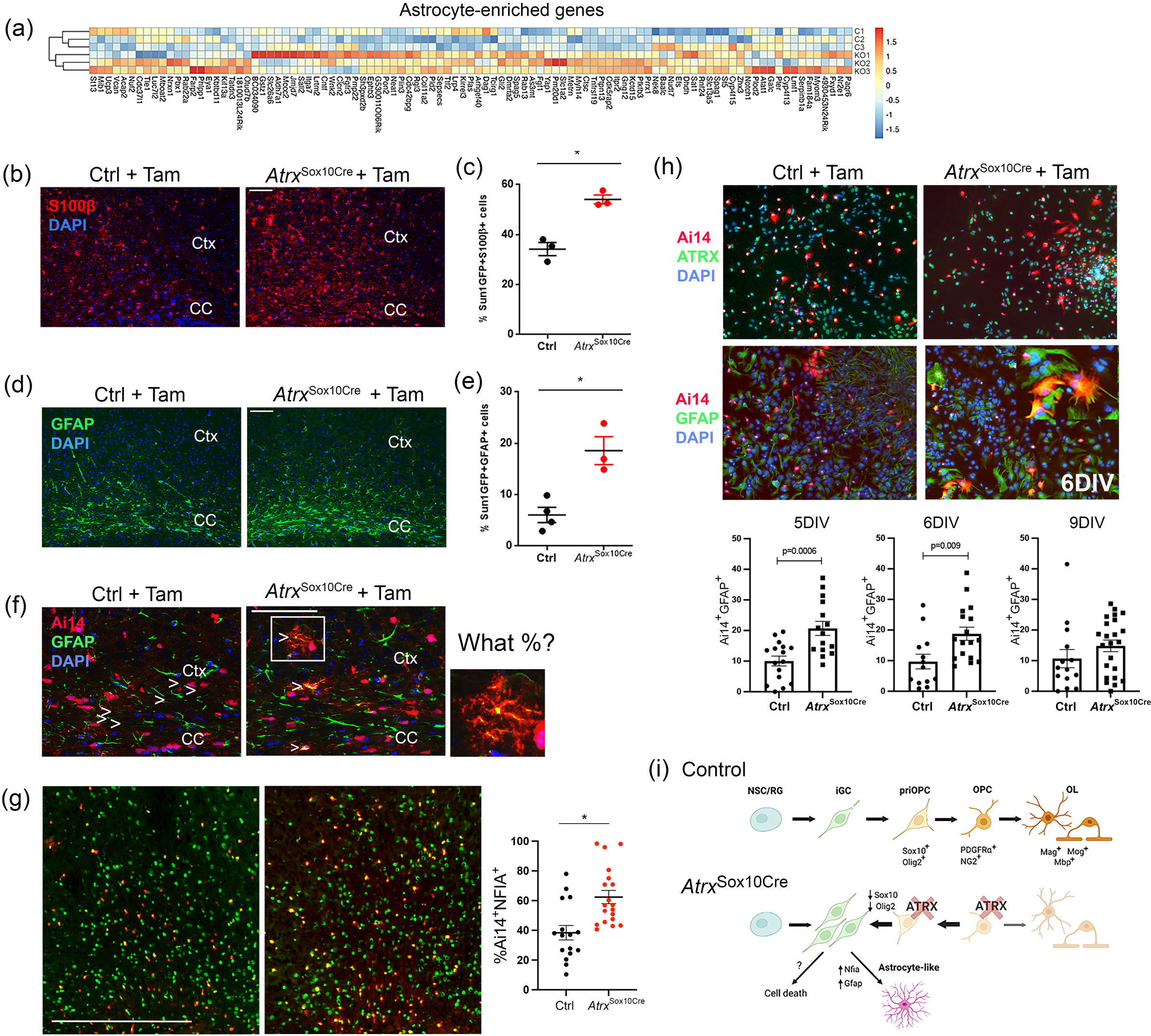
A subset of ATRX-null OPCs express astrocyte-enriched genes *in vivo* and *in vitro*. (a) Heatmap of astrocyte-enriched genes upregulated in ATRX-null cells. (b)lmmunofluorescence staining of the astrocyte marker S100β reveals increased levels In the P20 *Atrx*^sox10cre^ mouse corpus callosum (CC) and cortex (Ctx) compared to controls. Scale bar, 100 μm. (c) Quantification of the number of Sun1GFP^+^ OPCs that stain positive for S100β shows a significant overlap compared to control (n= 3-4 for each genotype). (d) lmmunofluorescence staining of the astrocyte marker GFAP shows increased staining in the P20 *Atrx*^sox1ocre^ mouse corpus callosum (CC) and cortex (Ctx) compared to controls. Scale bar, 100 μm. (e) Quantification of the number of Sun1GFP^+^ OPCs that stain positive for GFAP (n= 3-4 for each genotype). (f) lmmunofluorescence microscopy detects co-staining of the ere-sensitive Ai14 reporter (red) and the astrocyte marker GFAP (green) in the *Atrx*^sox10cre^ mouse brain *in vivo* (white arrowheads and higher magnification image on the right (g) lmmunofluorescence staining of NFIA (green) reveals increased expression in Ai14+ ATRX-null cells. Quantification is shown in graph on the right Scale bar, 400μm, (h) Overlap of Ai14 (red) and GFAP (green) in mixed glial cultures at 6DIV Quantification of Ai14+GFAP+ cells at 5, 6 and 9DIV (n=3). (i) Schematic representation of the effects of ATRX inactivation in Sox10+ pri-OPCs and OPCs in the postnatal forebrain. Loss of Olig2 and Sox10 expression reverts ATRX-mull OPCs and pri-OPCs to multipotent intermediate glial progenitor cells (iGCs), and gain the ability to differentiate into astrocytes in vitro and in vivo.. Created with BioRender.com. In all graphs, error bars represent +/−SEM and asterisks indicate significance (p<0,05 Student’s T-test)

Utilizing the Ai14 reporter to assess *Atrx*^Sox10Cre^ cell morphology, we could easily see that a subset of ATRX-null Ai14^+^ cells display a highly branched morphology typical of protoplasmic astrocytes as previously observed in *Olig2* knockout mice (*6, 7*). These cells co-expressed GFAP (**Figure 7f**) and the astrocyte specification factor NFIA (*92, 93*) (**Figure 7g**) indicating that a subset of ATRX-null cells gained the capacity to at least initiate astrocyte differentiation *in vivo*. We confirmed this finding in primary glial culture, where at 5 and 6 DIV, significantly more *Atrx*^Sox10Cre^ derived cells (Ai14^+^) expressed GFAP compared to control cells (**Figure 7h**). These results suggest that a portion of ATRX-null cells initiate an astrocytic differentiation program, instead of the Olig2-driven oligodendrocyte lineage pathway.

## Discussion

Chromatin remodelers have emerged as critical factors in the regulation of the OL lineage progression and myelination. Still, many questions remain as to the exact mechanisms by which they exert their effects. In the present study, we report key advances in our understanding of ATRX-mediated regulation of myelin development through both systemic and cell-autonomous mechanisms. The *Atrx*^Foxg1^Cre mice exhibit CNS hypomyelination that can partially be rescued by early postnatal administration of thyroid hormone. Cell type-specific inactivation of ATRX in OPCs, but not in neurons, also resulted in hypomyelination, showing that ATRX loss can have systemic as well as cell-autonomous roles in developmental myelination. We determined that ATRX directly binds to and activates the *Olig2* gene at regulatory sites and promotes an active chromatin state (H3K27Ac). Consequently, Olig2 protein level is reduced in a subset of ATRX-null OPCs, leading to a more permissive precursor state, perhaps through modified chromatin accessibility, allowing for ectopic astrogliogenesis.

We first demonstrated that myelination is defective in *Atrx*^FoxG1Cre^ mice, which have low levels of circulating T4 (*55, 79*). The thyroid hormone receptor is a transcriptional activator for several myelin genes, promoting differentiation of OPCs (*23–25*). We observed a significant yet partial rescue in myelinogenic genes and proteins in *Atrx*^FoxG1Cre^ mice following T4 supplementation. Whereas the number of APC-positive iOLs increased following T4 administration, the number of PDGFRα-positive OPCs was not restored to control levels. Furthermore, production of Olig2-expressing OPCs in *Atrx*^FoxG1Cre^ mice was not improved after thyroid hormone treatment, indicating that ATRX may be essential for OPC determination, proliferation or survival involving a mechanism distinct from T4 regulation.

Neuronal communication with OLs is emerging as a key regulator of myelination. Neurons have been proposed to be necessary for adaptive myelination (*94, 95*) and electrical activity from neurons has been shown to induce OPC proliferation and remyelination (*96*). ATRX is absent in all cell types of the forebrain of *Atrx*^FoxG1Cre^ mice, and it was thus conceivable that deletion of *Atrx* in neurons could cause a reduction of the OPC pool and subsequent myelination. Nonetheless, specific deletion of ATRX in neurons did not result in defects in the development of white matter. Conversely, deletion in Sox10-expressing pri-OPCs and OPCs had substantial deleterious effects on myelin production.

Our data indicate that in OPCs, ATRX binds several sites across the *Olig2* gene locus and promotes gene expression partly via H3K27Ac. Olig2 is a key determinant of OPC and OL fate, by promoting a cascade of gene expression, but also by directly suppressing astrocytic fate in conjunction with Olig1 (*3*). Loss of Olig2 can explain the reduced expression of Pdgfrα, since it is required for induction of this gene in cooperation with Sox10 (*8, 9*). Pdgfrα is essential for OPC proliferation and survival and the decreased expression of this gene might explain the reduced proliferation and increased cell death observed in *Atrx*^Sox10Cre^ mixed glial cultures. Depletion of the OPC pool can in turn cause the reduction in myelinating OLs and hypomyelination observed in *Atrx*^Sox10Cre^ mice.

The gliogenic switch observed upon loss of ATRX is most likely mediated by changes in the epigenetic and chromatin landscape. Two factors involved in this process are the histone deacetylase HDAC3 and the histone acetylase P300 (*83*). However, we found no evidence of overlap between the occupancy of ATRX and these two factors in OPCs. Rather, we discovered substantial enrichment of ATRX at H3K27Ac, CHD7 and CHD8 binding sites across the genome. Like ATRX, CHD7 and CHD8 are ATP-dependent chromatin-remodeling enzymes and they can bind to the enhancer regions and near transcription start sites of myelinogenic genes (*84, 86*). However, CHD8 expression appears to play a bigger role in OPCs than CHD7. Chromatin occupancy of CHD8 in OPCs is greater than that of CHD7 in OPCs (*97*), and CHD8 acts upstream of the Brg1-containing BAF complex and Olig2, both of which promote CHD7 expression (*86*). Moreover, CHD8 occupancy in OPCs corresponds to proximal enhancer regions marked by H3K27Ac, while this is not the case for CHD7. Based on this, we predict that ATRX co-binds chromatin sites with CHD8 (as opposed to CHD7) in OPCs, although the purpose of utilizing two chromatin remodelers at enhancers will need to be resolved in future studies. It is interesting to note that *CHD8* mutations cause congenital anomalies, intellectual disability and autism accompanied and severe white matter abnormalities (*98–100*), akin to ATR-X syndrome, suggesting that there might be mechanistic overlap in white matter pathogenesis caused by disruption of these chromatin remodelers.

The fate switch observed upon loss of ATRX is reminiscent of the effects of *Olig2* deletion in the mouse postnatal cortex (*6, 7*). The timing of *Olig2* ablation has been shown to reflect the number of OPCs that revert to an astrocytic fate. When *Olig2* is deleted constitutively in OPCs beginning at E16.5 there is almost complete conversion of these cells to astrocytes (*6*). However, deletion of Olig2 at P2 or P18 results in a 50% and 25% conversion to astrocytes, respectively (*7, 101*). From these results, we predict that an earlier inactivation of *Atrx* in OPCs might yield a greater conversion to the astrocyte lineage *in vivo*. Our *in vitro* system is designed for the mass production of OPCs and allows for more cell division than would normally be observed *in vivo*, especially considering that the proliferation capacity of OPCs declines with age (*102*). Accordingly, *Atrx*^Sox10Cre^ mixed glial cultures exhibit a more drastic increase in conversion to astrocytes over a period of 9DIV. Mechanistically, Olig2 has been reported to directly repress key astrocytic fate determinants, including NFIA and GFAP, both of which were inappropriately induced in ATRX-null OPCs. NFIA can convert primed chromatin to active chromatin during early stages of astrocyte differentiation (*92*), and has been shown to activate *Gfap* gene expression (*103*).

Overall, our study suggests that myelination deficits caused by ATRX deficiency stems from cell-intrinsic functions of ATRX in the oligodendrocyte lineage, in addition to systemic mechanisms. Our findings could have implications in the treatment of myelin deficits in patients with disruptive *ATRX* mutations. In addition to the potential benefits of thyroid hormone treatment, adjusting glial cell-fate imbalances and overcoming intrinsic defects in oligodendroglial cell maturation and the ensuing developmental dysmyelination could be important therapeutic targets to improve white matter defects in ATR-X syndrome patients.

## Author contributions

Rowland and Bérubé designed the study. Dumeaux analyzed the RNA-seq data, assisted in the ChIPseq analyses and reviewed the manuscript. Rowland, Jiang, Shafiq and Pena-Ortiz acquired the data. Rowland, Jiang, Ghahramani and Bérubé analyzed data and generated figures. Rowland and Bérubé wrote the manuscript.

## Acknowledgements

We thank Douglas Higgs and Richard Gibbons for kindly providing the *Atrx^loxP^* mice used in this study and Matt Edwards for generating the western blot in Figure 1e. Operating funds were from the Canadian Institutes for Health Research to NGB (MOP#142369). The authors have no conflict of interest. Please address all correspondence and requests for reprints to.

