## Supplementary files for "Systemic and intrinsic functions of ATRX in glial cell fate and CNS myelination"

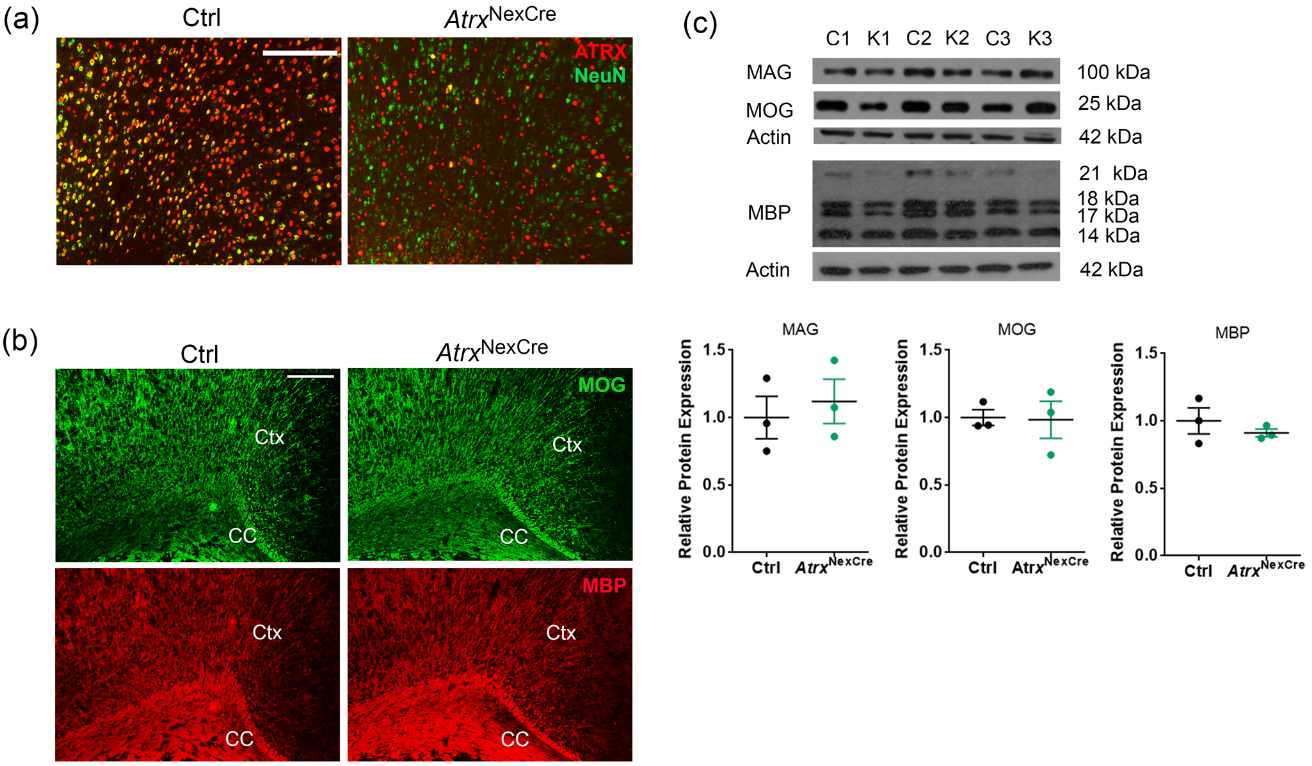


**Figure S1: Normal myelin levels in mice with *Atrx* deletion in forebrain excitatory neurons.** (a) Immunofluorescence staining of P20 brain cryosections shows absence of ATRX protein (red) in NeuN expressing neurons (green) in the cortex of *Atrx*$^{NexCre}$ mice. Scale bar, 100 µm. (b) MBP and MOG immunofluorescence staining of P20 *Atrx*$^{NexCre}$ and control brain cryosections. Scale bar, 100 µm. (c) Western blot analysis of MAG, MOG and MBP and quantification below shows similar levels of these proteins in the P20 *Atrx*$^{NexCre}$ mouse forebrain compared to controls when normalized to Actin protein levels (MAG p=0.63, MOG p=0.92, MBP p=0.43; Student's T-test). Error bars represent +/-SEM, n=3 each genotype.

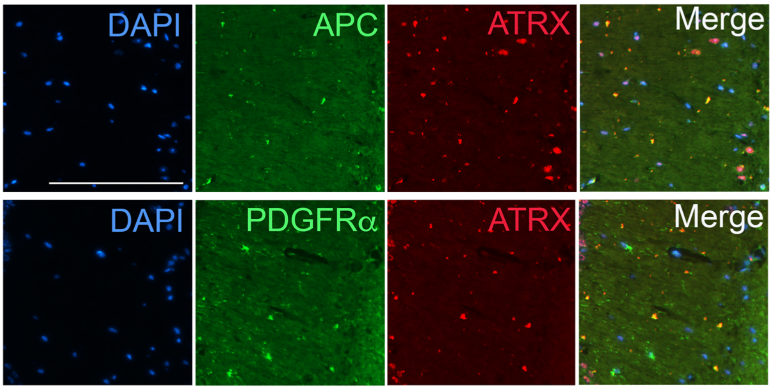


**Figure S2. ATRX protein is expressed in OPCs and OLs in the mouse brain.** Immunofluorescence staining of the mouse hippocampus with ATRX and PDGFRα (OPC marker). or ATRX and APC (OL marker)  Scale bar, 200μM.

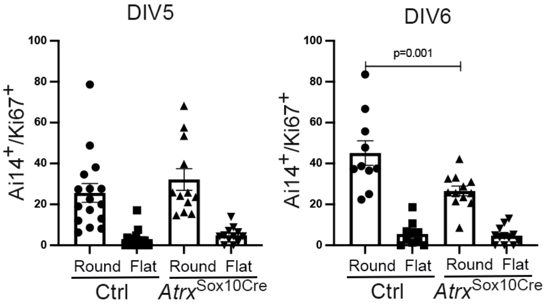


**Figure S3:** Ki67 staining reveals normal proliferation at DIV5, and decreased proliferation of round-shaped ATRX-null cells at DIV6. Error bars represent +/-SEM and asterisks indicate $p<0.05$ (Student's T-test).

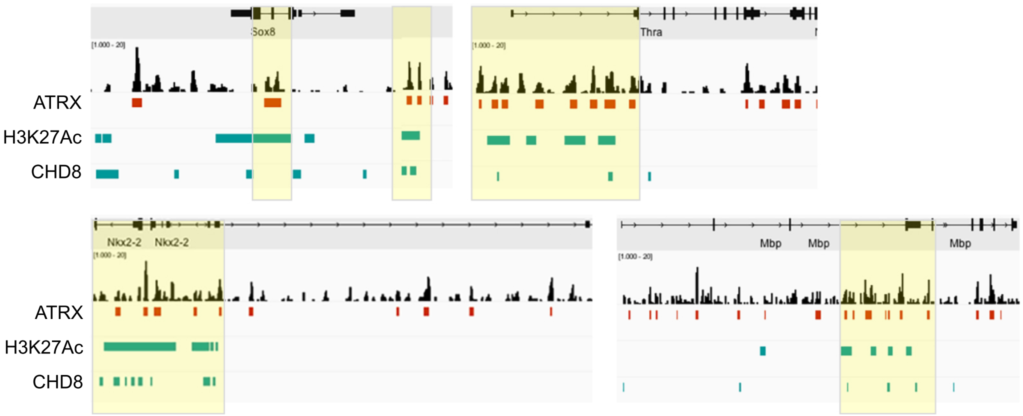


**Figure S4: ChIP-seq of ATRX, H3K27Ac and CHD8 at oligodendrocyte lineage genes in OPCs.**
UCSC views of genomic regions corresponding to *Sox8*, *Thra*, *Nkx2-2* and *Mbp* genes. ATRX ChIP-seq data is shown as sequence peaks in black and called peaks in red. H3K27Ac and CHD8 called peaks are shown in green. Yellow boxes highlight region of overlap.

### Supplementary Table 1: List of primer sequences

| Gene Name | Forward Primer | Reverse Primer | Application |
|---|---|---|---|
| AtrxWT | AGA AAT TGA GGA TGC TTC ACC | TGA ACC TGG GGA CTT CTT TG | Genotyping |
| Atrxfloxed | AGA AAT TGA GGA TGC TTC ACC | CCA CCA TGA TAT TCG GCA AG | Genotyping |
| FoxG1Cre | TGA CCA GAG TCA TCC TTA GCG | AAT GCT TCT GTC CGT TTG CC | Genotyping |
| NexCre | TGA CCA GAG TCA TCC TTA GCG | AAT GCT TCT GTC CGT TTG CC | Genotyping |
| Sox10Cre | CAC CTA GGG TCT GGC ATG T | CAG GTT TTG GTG CAC AGT CA | Genotyping |
| Sun1GFP | AAG GGA GCT GCA GTG GAG TA | CGG GCC ATT TAC CGT AAG TTA T | Genotyping |
| Ai14Tomato | GGC ATT AAA GCA GCG TAT CC | CTG TTC CTG TAC GGC ATG G | Genotyping |
| *Gapdh* | CAA CGA CCC CTT CAT TGA CCT | ATC CAC GAC CGA CAC ATT GG | qRT-PCR |
| *Atrx* | AGA AAT TGA GGA TGC TTC ACC | TGA ACC TGG GGA CTT CTT TG | qRT-PCR |
| *Olig2* | GCA GCG AGC ACC TCA AAT CT | AGA TCA TCG GGT TCT GGG GA | qRT-PCR |
| *Sox10* | CTA CAGG GAG TGC CCA CCT GG | GCT CTG TCT TTG GGG TGG TT | qRT-PCR |
| *Pdgfrα* | GTC AGG CCA CT AAA GAG GTC A | CGT GCA AGT TGA CAG CTT CC | qRT-PCR |
| *Olig1* | CTC GCC CAG GTG TTT TGT TG | CGA CGT GCC TTG CTA CCT AT | qRT-PCR |
| *Gpr17* | TCACAGCTTACCTGCTTCCC | TTGTCCGCATTGCTCAGAGT | qRT-PCR |
| *Cspg4* | CTCAAGATGGGAGCCTCAGC | ACAAAGGCGTCTGTCTGTGT | qRT-PCR |
| *Mag* | CAG AGA GCC ACT GCC TTC AA | TCA AAG GCC ACA GAG GTT C | qRT-PCR |
| *Mog* | AGA GGC AGC AAT GGA GTT GA | TGC GAT GAG AGT CAG CAC AC | qRT-PCR |
| *Mbp* | CAT TGG GTC GCC ATG GGA AA | AGC CTC TCC TCG GTG AAT CT | qRT-PCR |
| *Plp* | GGC CAC TGG ATT GTG TTT CT | GAA AGC ATT CCA TGG GAG AA | qRT-PCR |
| *Gfap* | AAA CCA GCC TGG ACA CCA AA | ACA CCT CAC ATC ACC ACG TC | qRT-PCR |
| *S100β* | TGA AGC CAG AGA GGA CTC CA | CCA GGA AGT GAG AGA GCT CG | qRT-PCR |
| Olig2A | ATT CCC CGT CTC ACT CCG TA | TAG CGG GGC TGC TAA AGA AG | ChIP qRT-PCR |
| Olig2B | GCC TGA CGC TAC AGT GAC AA | TAG CGG GGC TGC TAA AGA AG | ChIP qRT-PCR |
| Olig2C | CTG CCT CCA CCC AGC TA TAA | GGT GTT GGC TCG GTC TGT AA | ChIP qRT-PCR |
| Olig2D | ATC TTC CTC CAG CAC CTC CT | GTT CGC GGC TGT TGA TCT TC | ChIP qRT-PCR |
| Olig2E | TAG GAG ACT CCC AGG AAC CG | GAC CAC AGG ACC CTA AGT GC | ChIP qRT-PCR |
| Olig16kb | CCT CCC ACA CAC ACA CCT TT | AGG TGG AAG GTT AGG AGG CA | ChIP qRT-PCR |
